# Phollow: Visualizing Gut Bacteriophage Transmission within Microbial Communities and Living Animals

**DOI:** 10.1101/2024.06.12.598711

**Authors:** Lizett Ortiz de Ora, Elizabeth T Wiles, Mirjam Zünd, Maria S Bañuelos, Nancy Haro-Ramirez, Diana S Suder, Naveena Ujagar, Julio Ayala Angulo, Calvin Trinh, Courtney Knitter, Shane Gonen, Dequina A Nicholas, Travis J Wiles

## Abstract

Bacterial viruses (known as “phages”) shape the ecology and evolution of microbial communities, making them promising targets for microbiome engineering. However, knowledge of phage biology is constrained because it remains difficult to study phage transmission dynamics within multi-member communities and living animal hosts. We therefore created “Phollow”: a live imaging-based approach for tracking phage replication and spread in situ with single-virion resolution. Combining Phollow with optically transparent zebrafish enabled us to directly visualize phage outbreaks within the vertebrate gut. We observed that virions can be rapidly taken up by intestinal tissues, including by enteroendocrine cells, and quickly disseminate to extraintestinal sites, including the liver and brain. Moreover, antibiotics trigger waves of interbacterial transmission leading to sudden shifts in spatial organization and composition of defined gut communities. Phollow ultimately empowers multiscale investigations connecting phage transmission to transkingdom interactions that have the potential to open new avenues for viral-based microbiome therapies.

## INTRODUCTION

Bacteriophages (or “phages”) are viruses that infect bacteria. As the most numerous biological entities on Earth, phages exert tremendous influence over microbial communities^1–3^. Phages are largely recognized as lethal predators of bacteria but they can also enhance bacterial fitness by mediating the transfer of novel genetic traits^4–6^. Identifying how phages shape the form and function of microbiomes could inspire approaches for improving human and environmental health^7–10^. For example, controlled phage outbreaks could be used to deplete disease-causing pathobionts or encourage the spread of beneficial activities. Ultimately, however, harnessing phages to kill or cultivate bacterial communities requires a contextual understanding of phage replication regimes.

Lytic phage replication is a form of horizontal transmission that is overtly antagonistic; phages infect and take over the machinery of bacterial cells to produce new virions and often disperse through the explosive lysis and death of their host. In contrast, lysogenic replication is a form of vertical transmission that is frequently associated with mutualistic interactions^5,6^. During lysogenic replication, phage genomes are maintained as integrated or episomal prophages that replicate with the bacterial chromosome. Phages capable of forming lysogenic partnerships are referred to as temperate phages.

Through the lens of microbiome engineering, phage replication regimes can be viewed as potential targets for manipulating resident microbial communities. However, the factors governing phage replication and transmission in situ are largely unknown^2,11–13^. This is especially the case within the confines of the gastrointestinal tracts of humans and other animals, where many questions about phage biology remain unanswered. Are outbreaks of lytic replication spatially widespread or locally restricted? Do they occur over rapid or short timescales? What are the consequences of phage replication dynamics on the broader bacterial community, and how might they affect cells and tissues of the animal host? These questions remain largely unexplored because conventional approaches lack spatiotemporal sensitivity and scalable resolution^11,12^. Without knowing the location and duration of phage replication or whether phages are in the form of extracellular virions or intracellular prophages, building a contextualized understanding of phage biology within the gut is severely constrained.

In this paper we overcome current limitations by creating “Phollow”: a collection of tools and techniques for monitoring phage outbreaks in situ by live imaging. We demonstrate that Phollow enables the multiscale tracking of phages in both their viral and cellular forms, and their interactions with bacteria and animal tissues. Combining Phollow with optically transparent zebrafish revealed that virions can be taken up by cells lining the gut, including enteroendocrine cells, and that they quickly disperse to extraintestinal tissues including the liver and brain as well as to the environment outside the animal host. We also found that abrupt changes in phage replication regimes lead to rapid shifts in community spatial organization and composition. Altogether, we show that Phollow is a tractable solution to investigating phage transmission dynamics in the context of microbial communities and living animals.

## RESULTS

### Model phage selection

We selected P2-like phages as a model system for developing approaches to visualize phage replication regimes. P2-like phages belong to the Caudoviricetes class of tailed phages and infect over 120 bacterial genera across the phylum Proteobacteria^14–16^. P2-like phages display a temperate lifestyle, which enables the study of both lytic and lysogenic replication. Lytic replication of the P2-like phages used in our study is induced by DNA damage and activation of the “SOS” response (Extended Data Fig. 1). Although P2-like phage lineages have been extensively studied, little is known about their in situ replication and transmission dynamics.

### Design, construction, and infectivity of Phollow phages

We devised an in vivo tagging system in which phage virions are fluorescently marked during intracellular assembly, but then become marked by a different color upon infection and replication within a new bacterial host. This combinatorial labeling strategy—which we call “Phollow”—makes it possible to *follow* chains of interbacterial transmission during phage outbreaks.

We constructed fluorescently marked “Phollow phages” using the SpyTag:SpyCatcher tagging system^17,18^ (Fig. 1a and Extended Data Table 1). The major capsid protein (GpN) of a P2-like prophage harbored within *Escherichia coli* HS^19,20^ (a human gut-derived commensal) was modified to contain a C-terminal peptide linker and SpyTag. We inserted a Tn*7* transposon into the bacterial chromosome carrying a constitutively expressed gene encoding SpyCatcher fused to one of several spectrally distinct fluorescent proteins. SpyTags decorating P2 capsids become covalently bound by SpyCatcher proteins during virion assembly, generating fluorescently marked Phollow phages (Fig. 1a). We refer to engineered bacterial strains harboring Phollow prophages as “Phollow virocells”. We note that we also assessed the functionality of other SpyTag variants^21^ and the SnoopTag:SnoopCatcher system^22^ (key observations are summarized in Extended Data Fig. 2 and Extended Data Table 2).

**Figure 1.**
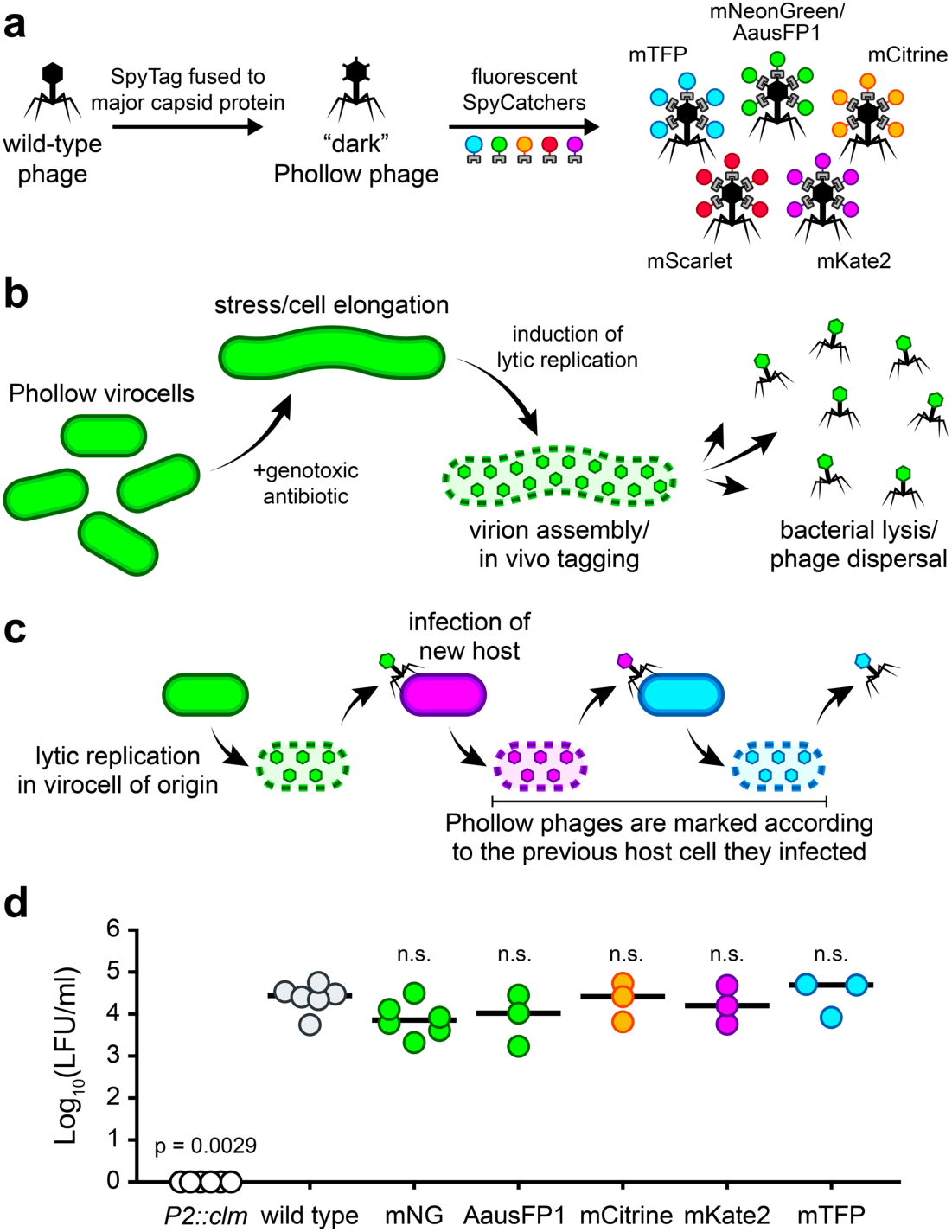
Design, construction, and infectivity of Phollow phages. **(a)** Design overview of P2 Phollow phages using the SpyTag:SpyCatcher system. The major capsid protein (GpN) of temperate phage P2 is first modified to contain a C-terminal SpyTag (creating a “dark” Phollow phage). Next, a SpyCatcher peptide, containing a C-terminal fusion to one of several fluorescent proteins, is constitutively expressed in trans from the bacterial chromosome to produce differentially tagged Phollow phages. **(b)** Cartoon schematic of P2 phage-tagging in vivo using Phollow. First, Phollow virocells (i.e., bacterial cells carrying a SpyTag-modified P2 prophage and expressing a fluorescent SpyCatcher protein) are treated with a genotoxic antibiotic to induce lytic replication of the P2 Phollow phage. Phollow virocells display hallmark morphological changes upon activation of the DNA damage “SOS” response; namely, cell filamentation. As P2 phage gene activation and replication proceed, the fluorescent SpyCatcher protein redistributes from the cytosol to sites of capsid assembly. Upon bacterial cell lysis, fluorescently tagged P2 Phollow phage virions are released. **(c)** Cartoon schematic of combinatorial Phollow tagging to track interbacterial phage transmission. Shown is an mNeonGreen Phollow phage (green) lytically replicating and dispersing to infect a susceptible target bacterial cell expressing an mKate2 SpyCatcher peptide (magenta). Subsequent lytic replication in the mKate2 target host cell produces mKate2 Phollow phages that then spread to infect a susceptible target bacterial cell expressing an mTFP SpyCatcher peptide (cyan). Lytic replication in the mTFP target cell host then produces mTFP Phollow phages. **(d)** Phollow phages display similar levels of infectivity compared to a wild-type P2 phage. The infectivity of wild-type and differentially marked Phollow phages was assessed by measuring lysogen-forming units (LFU) generated from infection of an *E. coli* HS target cell cured of its resident P2 prophage. Lysogeny was monitored using prophages carrying a chloramphenicol resistance gene. *P2::clm* is an *E. coli* HS strain that carries a chloramphenicol resistance marker in place of the P2 prophage genome and is used to control for phage-independent transfer of chloramphenicol resistance. Note: we found that mScarlet Phollow phages were technically difficult to image and therefore did not characterize their infectivity. Bars denote medians and each circle represents an independent biological replicate. Significant differences compared to “wild type” were determined by one-way ANOVA using a Kruskal-Wallis and Dunn’s multiple comparisons test (p < 0.05; n.s. = not significant).

As designed, SpyCatcher proteins redistribute from the bacterial cytosol to assembling Phollow phage capsids during lytic replication (Fig. 1b). Host lysis then leads to the dispersal of fluorescently marked Phollow phages throughout the environment (Fig. 1b). Tracking interbacterial transmission is accomplished using target bacterial cells expressing fluorescent SpyCatcher variants that are a different color than those expressed by the original Phollow virocell. Infection of differentially marked bacterial hosts, therefore, produces Phollow phages donning new fluorescent tags (Fig. 1c). We confirmed that Phollow phages exhibit similar infectivity compared to a non-fluorescently tagged phage, which emphasizes their capacity for recapitulating wild-type replication and transmission (Fig. 1d). Below, we characterize Phollow as an experimental approach and demonstrate its utility by investigating unexplored aspects of P2 phage biology.

### Investigating the cell biology of phage lytic replication

We first applied Phollow to probe intracellular features of P2 phage lytic replication. Remarkably, we captured the complete sequence of Phollow phage assembly and dispersal within a single field of view (Fig. 2a). Treating *E. coli* Phollow virocells with the DNA-damaging agent mitomycin C (MMC) led to cellular filamentation and the formation of fluorescent viral foci, culminating in bacterial lysis and the extracellular release of virions. Leveraging the cell virological traits of lytically replicating Phollow phages, we used imaging flow cytometry to quantify MMC induction kinetics. We collected cells at regular intervals spanning initial treatment to bacterial lysis (Fig. 2b). Gating on features of cell filamentation and the presence of viral foci (Fig. 2c), we found that peak induction occurred 1h post-MMC treatment and comprised ∼20% of the bacterial population (Fig. 2d). The number of cells with lytically replicating phage fell sharply over time, corresponding to cell lysis (Fig. 2b,d).

**Figure 2.**
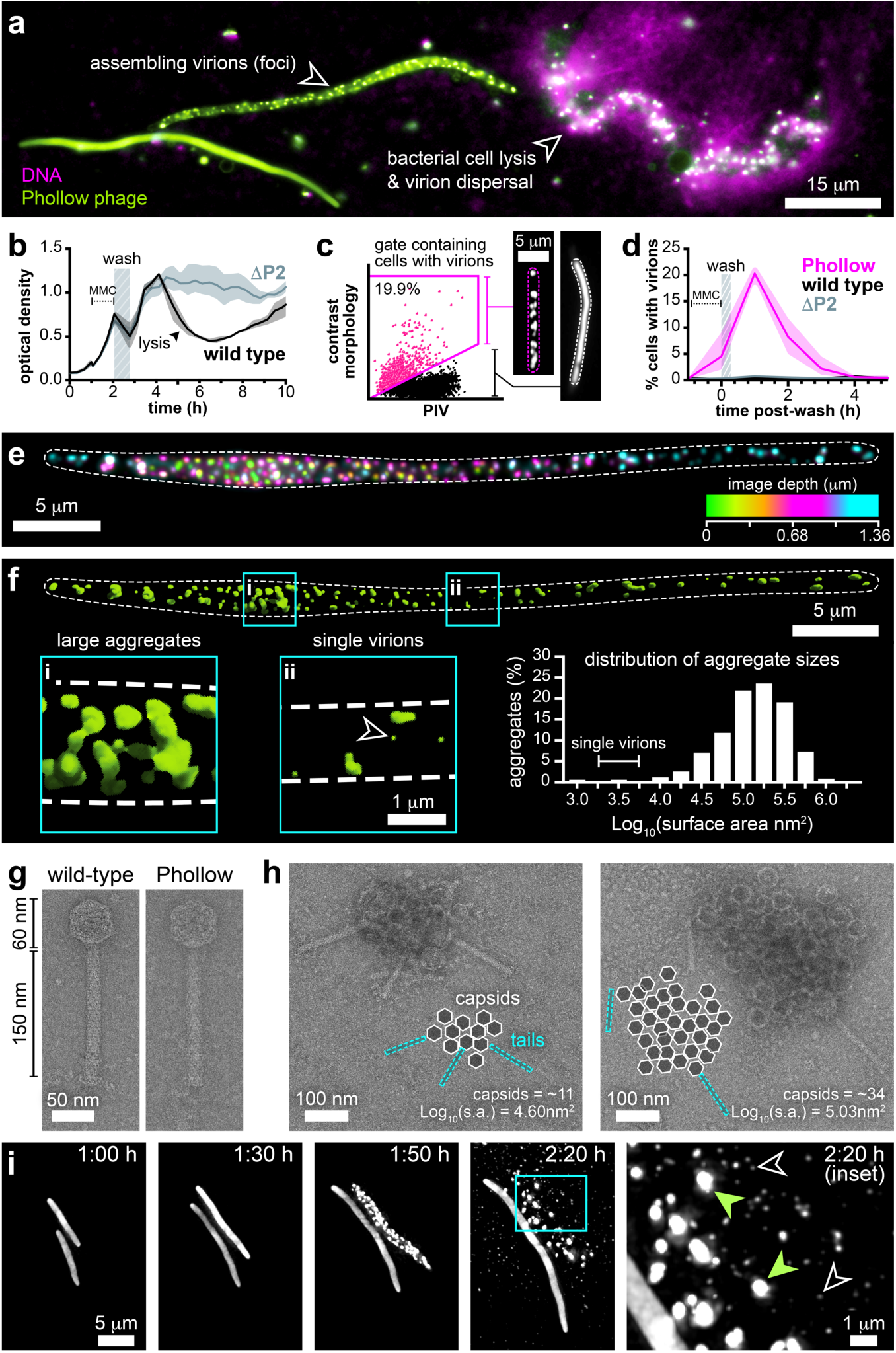
Investigating the cell biology of phage lytic replication. **(a)** Single field of view showing three mNeonGreen Phollow virocells (derived from *E. coli*) in different phases of lytic replication induced by MMC. (left) A cell undergoing filamentation. (middle) A filamented cell containing assembling virions. (right) A cell undergoing lysis and release of viral particles. DNA from the lysing cell is labeled with the cell-impermeable DNA dye EthD-III (magenta). **(b)** Representative induction (or “lysis”) curve of P2 phage lytic replication in *E. coli* HS. The optical density of cultures (at 600 nm) containing either wild-type (black line) or a P2 prophage-cured strain (ΔP2, gray line) was monitored prior to and following a 1h MMC treatment. Shaded regions represent the range of optical density measurements from three technical replicates. MMC was washed from cultures at 2h (hashed bar), which prevents interference of phage lytic replication. Washing also produces a slight decrease in cell density due to some cell loss. An abrupt drop in optical density of the wild-type culture at 4h marks widespread lysis, which is not observed in the ΔP2 culture. **(c)** Shown is an example imaging flow cytometry gating scheme used to quantify cells harboring lytically replicating phages (marked by magenta box and hashed line). Sample is taken from a culture at peak lytic replication (∼1h post-MMC treatment). PIV = pixel intensity variance. **(d)** Quantification of lytic replication over time following MMC treatment using imaging flow cytometry as in panel c. Cultures were treated and washed as in panel b. Phollow virocell cultures (magenta, “Phollow”) produce a sharp peak of cells with lytically replicating phage at ∼1h post-MMC treatment. “Wild type” (black) and “ΔP2” (gray) cultures serve as controls and do not produce cells harboring viral foci. Shaded regions represent ranges from three biological replicates. **(e)** Z-projected image of a bacterial cell harboring viral foci. Image represents 1.36 μm of total sample depth (16 images captured using 0.085 μm steps). Z-planes are pseudo-colored according to z-depth to highlight the scattered distribution of viral foci throughout the cell. White dashed line marks the cell perimeter. **(f)** 3D render of the bacterial cell shown in panel e, highlighting that viral foci are likely virion aggregates. Inset “i” shows an area of the cell with large virion aggregates. Inset “ii” shows an area of the cell containing single virions (arrowhead). Histogram in the bottom right shows the distribution of aggregate sizes based on their 2D surface area (n = 357 aggregates from across 5 individual cells). A single P2 capsid is expected to have a diameter of 50– 60 nm, producing a 2D Log_10_(surface area) of 3.29–3.45 nm^2^. **(g)** Representative TEM micrographs of an unmodified wild-type (left) and an mNeonGreen Phollow phage (right). **(h)** TEM micrographs of two example extracellular virion aggregates of unmodified wild-type P2 phages. Inset cartoons diagram the relative distribution of capsid and tail components within each aggregate. The surface area (s.a.) of each aggregate was estimated based on the area encompassing discernable capsid structures. **(i)** Single frames extracted from Supplementary Movie 2 showing the induction, assembly, and dispersal of virion aggregates. Upon cell lysis at 2:20h (inset), single and small virion aggregates rapidly diffuse throughout the extracellular environment (black arrowheads) whereas large virion aggregates remain at the site of lysis (green arrowheads).

We next inspected the intracellular assembly and organization of Phollow phage virions. At peak lytic replication, we estimate that Phollow virocells harbor an average of 1.6 ± 0.4 viral foci per micron cell length (mean length = 44.92 μm ± 7.11, n = 5) (Fig. 2e). Viral foci most often displayed a scattered distribution throughout the cell; however, we also observed serpentine patterns that could indicate early organizational dynamics during virion assembly (Supplementary Movie 1, and Extended Data Fig. 3). Analyzing 3D projections of viral foci further revealed that they have an average surface area that is ∼100 times larger than what is expected for a single P2-like phage capsid with a diameter of 60 nm^14^, suggesting that they represent multi-virion aggregates (Fig. 2f). To determine whether virion aggregates are a consequence of Phollow tagging, we compared unmodified wild-type and Phollow phage virions by transmission electron microscopy (TEM). There was no evidence of overt structural differences between the two phages (Fig. 2g, and Extended Data Fig. 4a). Notably, TEM revealed aggregates of wild-type P2 phages that were similar in size to those identified by fluorescence microscopy (Fig. 2h, and Extended Data Fig. 4b). The aggregates we observed contained ∼10–40 viral capsids with the occasional presence of tail tubes facing outward in a “pinwheel-like” formation (Fig. 2h).

To ascertain the fate of virion aggregates, we performed time lapse imaging of Phollow phage assembly and dispersal. Phollow virocells were imaged while growing on agar pads containing MMC. At approximately 2h, aggregates began to form within cells, followed by widespread bacterial lysis 20–30 minutes later (Fig 2i and Supplementary Movie 2). Aggregates quickly dispersed into clouds of rapidly diffusing particles, suggesting that bacterial lysis and viral disaggregation are coordinated (Fig. 2i, 2:20h inset). Altogether, these experiments demonstrate the utility of Phollow for studies of cellular virology, from intracellular induction and virion assembly to bacterial lysis and dispersal.

### Monitoring virion dispersal by fluorescence microscopy and flow virometry

Quantifying extracellular virions can be used to estimate the potential for horizontal transmission. However, differentiating viral particles from cellular debris and vesicles is challenging with high-sensitivity imaging and flow cytometry techniques^23^. Underscoring this problem, many vesicles produced during lytic replication contain cytosolic contents (including virions) and DNA (Fig. 3a,b). We found that implementing established purification steps^24^ facilitated straightforward identification of virions by microscopy (Fig. 3b, bottom) but that the residual debris in viral preparations continued to interfere with flow cytometric analysis. We therefore utilized Phollow virocell control strains to develop a flow virometry gating strategy that could differentiate free viral particles from cell debris and thus, enable their quantification. First, we discerned small particles and cell debris originating from bacteria expressing a fluorescent SpyCatcher protein but not carrying a P2 prophage (Fig. 3c, left). We then identified fluorescent debris and vesicles associated with lytic replication using bacteria expressing a fluorescent SpyCatcher protein and carrying an unmodified wild-type prophage (Fig. 3c, middle). With these gates established, we could confidently quantify fluorescently marked Phollow phages (Fig. 3c, right). We further found that similar custom gating strategies can be used for fluorescent colors aside from mNeonGreen (Fig. 3d), and that it is possible to parse mixed populations of differentially marked Phollow phages (Fig. 3e).

**Figure 3.**
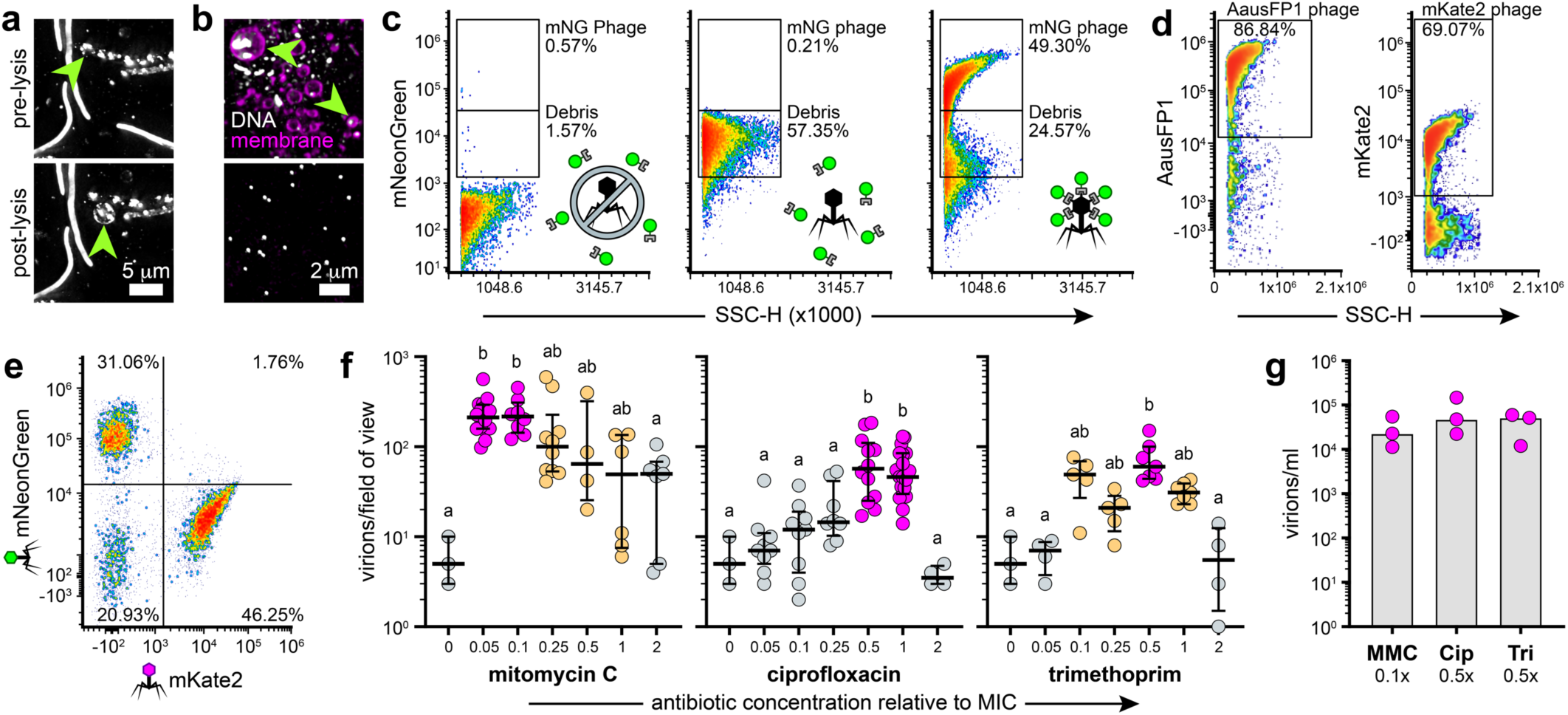
Monitoring virion dispersal by fluorescence microscopy and flow virometry. **(a)** Maximum intensity projection image of MMC-induced mNeonGreen Phollow virocells before (top) and after a lysis event (bottom). Green arrowheads indicate a cell that gives rise to a membrane vesicle containing virion aggregates and cytosolic SpyCatcher protein. **(b)** Top: MMC-induced cultures of *E. coli* carrying “dark” Phollow phages. Staining the culture with the membrane dye FM 4-64 and DNA dye Hoechst 33342 reveals that numerous membrane vesicles are generated during cell lysis, which frequently contain DNA (green arrowheads). Bottom: Purified lysates show dramatically reduced levels of membrane vesicles and regularly sized DNA-positive puncta that are likely individual virions. **(c)** Representative flow virometry plots showing the gating strategy for quantifying Phollow phage virions in purified lysates. Left: A MMC-treated *E. coli* HS population cured of their P2 prophage and expressing an mNeonGreen (mNG) catcher peptide establishes a baseline for fluorescent viral-like particles. Middle: A MMC-treated *E. coli* HS population carrying a wild-type P2 prophage and expressing an mNG catcher peptide is used to establish a generic “Debris” gate resulting from phage-driven cellular lysis. Right: A MMC-treated *E. coli* HS mNG Phollow virocell population is used to quantify production of fluorescently marked viral-like particles. **(d)** Representative flow virometry plots showing gates for AausFP1 (left) and mKate2 (right) Phollow phage virions. Gates were set as in panel c but “Debris” gates are not shown. **(e)** Representative flow virometry plot showing the quantification of Phollow phage virions from a mixed lysate. **(f)** Fluorescence microscopy-based quantification of Phollow phage virion production in response to treatment with the genotoxic antibiotics mitomycin C (left), ciprofloxacin (middle), or trimethoprim (right). Antibiotic concentrations are given relative to each antibiotic’s minimum inhibitory concentration (MIC) against wild-type *E. coli* HS. Each circle represents a distinct and non-overlapping field of view; data were pooled from 3 biological replicates. Data for the “0” concentration is the same for all plots and is used to set a statistical baseline. Statistical groupings (denoted by letters and color-coding) in each plot were determined using one-way ANOVA with a Kruskal-Wallis and Dunn’s multiple comparisons test (p < 0.05). **(g)** Flow virometry-based quantification of Phollow phage virion production induced by mitomycin C (MMC, 0.1x MIC), ciprofloxacin (Cip, 0.5x MIC), or trimethoprim (Tri, 0.5x MIC). Bars indicate medians derived from three biological replicates (circles). No statistical differences were found by one-way ANOVA using a Kruskal-Wallis and Dunn’s multiple comparisons test (p < 0.05).

To demonstrate how Phollow enables imaging and flow virometry-based studies, we compared the magnitude of virion dispersal induced by three different genotoxic antibiotics. We asked this question to clarify how genotoxic antibiotics with different modes of action and minimum inhibitory concentrations (MIC) may impact the potential for horizontal transmission (Extended Data Fig. 5). Screening virion production by fluorescence microscopy (Fig. 3f) first revealed that MMC, a DNA crosslinking agent, displays potent activity over a broad concentration range, with peak virion output occurring at 0.1x MIC. In contrast, the DNA gyrase inhibitor ciprofloxacin induces the production of viral particles across a narrow concentration range, peaking between 0.5–1x MIC. Lastly, trimethoprim, which interferes with thymidine synthesis, induces virion production over a slightly broader concentration range compared to ciprofloxacin, peaking at 0.5x MIC. Using flow virometry, we found that peak virion output across the three antibiotics was essentially equal, suggesting that they have similar capacities for spurring lytic replication despite their unique induction profiles (Fig. 3g). Together, this initial survey of antibiotic-specific induction activities demonstrates how Phollow expedites imaging and flow virometry-based studies capable of uncovering contextual modulators of virion dispersal.

### Visualizing outbreaks of phage lytic replication within the vertebrate gut

We next combined Phollow with optically transparent larval zebrafish, which enable the three-dimensional organization of microbial communities throughout a vertebrate intestine to be captured by live imaging^25–27^ (Fig 4a and Supplementary Movie 3). The small size of larval zebrafish and the aqueous media in which they live also make it possible to track virion dispersal across the entire body and the external environment.

**Figure 4.**
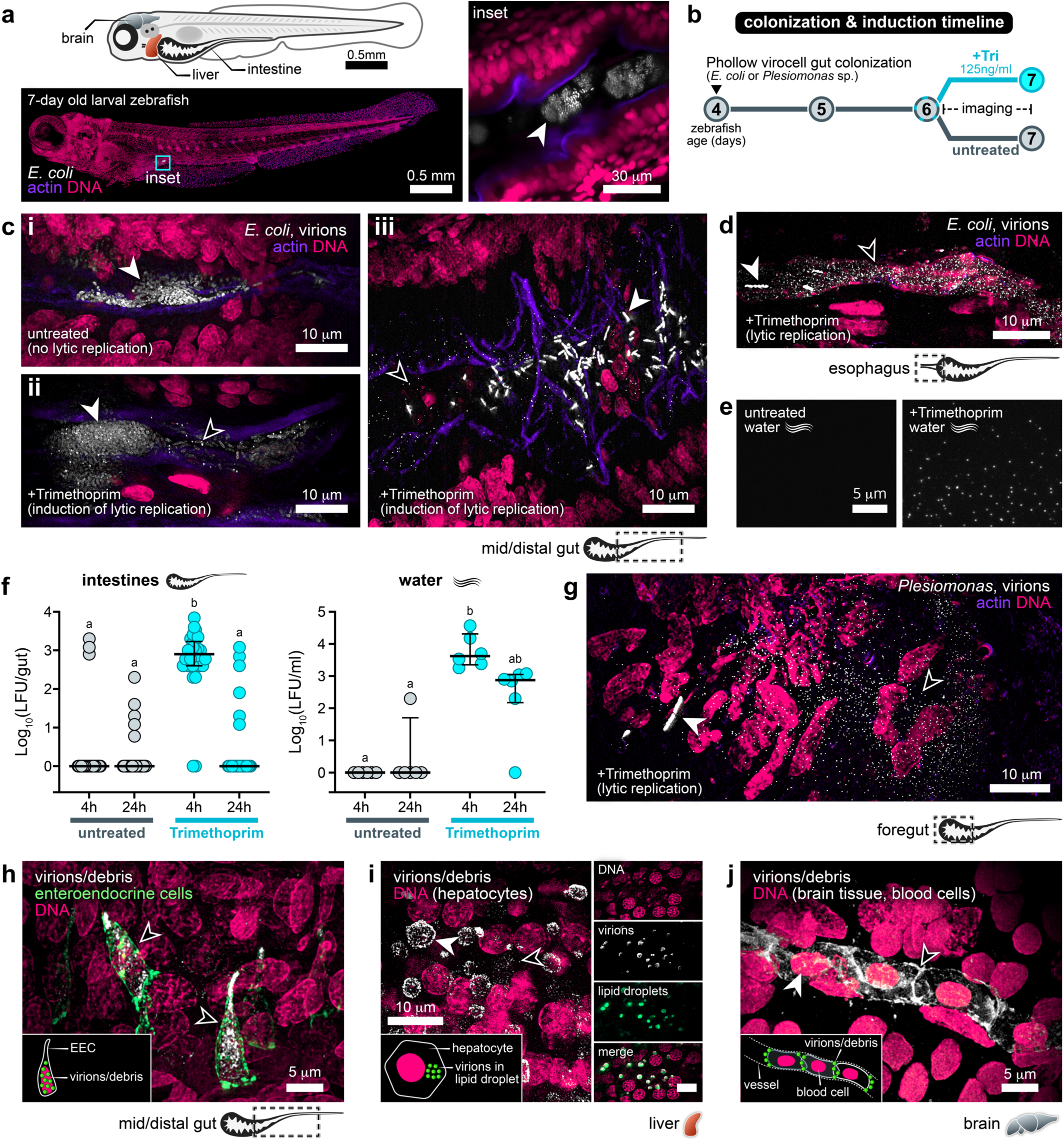
Visualizing outbreaks of phage lytic replication within the vertebrate gut. **(a)** Top: Schematic of a larval zebrafish with relevant anatomical features highlighted. Bottom: Maximum intensity projection of a 7-day old larval zebrafish colonized by *E. coli* generated from 297 tile-scanned and merged images acquired by super-resolution microscopy. (right) Inset is a single image of an intestinal region containing luminal aggregates of *E. coli* (arrowhead). DNA (magenta, JF549 dye) and actin (purple, CF405-phalloidin) highlight intestinal structure. **(b)** Diagram of bacterial colonization and antibiotic induction timeline. “Tri” = Trimethoprim. **(c)** (i) Maximum intensity projection image of an untreated *E. coli* Phollow virocell population within the gut (white arrowhead). (ii, iii) Two example maximum intensity projection images of trimethoprim-treated *E. coli* Phollow virocell populations. White arrowheads mark bacterial aggregates and single cells; black arrowheads mark viral particles. Images were acquired 4h post-treatment. **(d)** Maximum intensity projection image showing viral particles in the esophageal region. White arrowhead marks a single cell; black arrowhead marks viral particles. Image was acquired 4h post-treatment. **(e)** Fluorescence microscopy images of water samples taken from untreated (left) or trimethoprim-treated (right) zebrafish. An abundance of viral particles is evident in treated samples. Images were acquired 4h post-treatment. **(f)** Quantification of infectious virions from dissected intestinal tissues (left) and water (right) at 4h and 24h post-trimethoprim treatment (cyan circles). Quantification of infectious virions from time-matched untreated samples are shown in gray. Infectious virions are estimated by measuring the number of lysogen-forming units (LFU) per gut or ml of water. Statistical groupings (denoted by letters) in each plot were determined using one-way ANOVA with a Kruskal-Wallis and Dunn’s multiple comparisons test (p < 0.05). **(g)** Maximum intensity projection image showing viral particles derived from *Plesiomonas* Phollow virocells within the intestine. White arrowhead marks a single cell; black arrowhead marks viral particles. Image was acquired 4h post-treatment. **(h)** Maximum intensity projection image showing viral particles derived from *Plesiomonas* Phollow virocells within enteroendocrine cells (EEC, green). The membranes of EECs are marked using TgBAC(*nkx2.2a*:*megfp*) reporter fish. Black arrowheads mark EECs containing viral particles or phage debris. Inset shows a cartoon representation of viral particles associating with an EEC. Image was acquired 4h post-treatment. **(i)** Left: Maximum intensity projection image showing viral particles derived from *Plesiomonas* Phollow virocells associating with hepatocytes of the liver. Hepatocytes can be disguised based on the circular morphology of their nuclei^29^. White arrowhead marks virions within a vesicle-like structure; black arrowhead marks a virion aggregate or single phage. Inset shows a cartoon representation of viral particles associating with hepatocytes. Right: montage shows liver tissue from a separate fish stained with BDP (green) to mark lipid droplets. Image was acquired 4h post-treatment. **(j)** Maximum intensity projection image showing viral particles derived from *Plesiomonas* Phollow virocells within a blood vessel in the brain. White arrowhead marks a nucleated red blood cell; black arrowhead marks virions or phage debris associating with the surface of blood cells. Note: the opacity of brain tissue made it difficult to determine the presence of individual viral particles in this sample. Inset shows a cartoon representation of viral particles associating with red blood cells. Image was acquired 4h post-treatment.

We colonized the intestines of four-day old germ-free larval zebrafish with AausFP1 *E. coli* Phollow virocells, and after two days, fish were treated for 24h with trimethoprim or left untreated (Fig. 4b). AausFP1 was used for in vivo imaging experiments because it is one of the brightest known green fluorescent proteins^28^. We chose trimethoprim because it induces lytic replication over a relatively broad concentration range and it likely has fewer adverse effects on animal physiology compared to MMC. Within 4h of administering trimethoprim, clouds of viral particles manifested throughout the gut, from the foregut (including the esophagus) to the distal gut (Fig. 4c,d). Outbreaks of lytic replication were acute and tapered off starting around 8h post-treatment with little trace of viral particles after 24h. Notably, most untreated animals did not harbor viral particles but in some instances we could observe spontaneous lytic replication (Fig. 4c inset ‘i’, Extended Data Fig. 6a). Trimethoprim-induced lytic replication within the gut was always accompanied by the emergence of virions in the external water environment (Fig. 4e). The dispersal of viral particles throughout the water appears to be driven by their expulsion from the intestine since we did not find evidence of virion production in the absence of animal hosts (Extended Data Fig. 6b).

We next determined if the patterns of Phollow phage outbreaks revealed by live imaging were mirrored by fluctuations in numbers of infectious virions. We found that lysogen-forming units (LFUs) from dissected gut tissues were only sporadically detected in untreated animals; however, after 4h of trimethoprim treatment the majority of samples contained high numbers of LFUs (Fig. 4f, left). By 24h post-treatment, LFUs returned to baseline. Within the water, LFUs generally followed trends in the intestine with two notable differences (Fig. 4f, right). First, in untreated water samples, the frequency of detecting LFUs was incredibly low despite spontaneous induction in some animals. Second, the abundance of LFUs in treated water samples was relatively sustained over the 24h sampling period. These observations together suggest that, following trimethoprim treatment, viral particles are continuously expelled from the gut and accumulate in the water where they display a moderate rate of inactivation.

Our findings motivated us to further probe whether related phages from diverse bacterial hosts display unique or overlapping behaviors within the gut. Within our collection of zebrafish gut bacterial isolates, we identified a functional P2-like prophage harbored by a strain of *Plesiomonas*^30,31^. *Plesiomonas* species are frequently isolated from the microbiomes of aquatic animals and can cause gastroenteritis in humans^32^. We found that *Plesiomonas* Phollow virocells displayed patterns of intracellular virion assembly and extracellular dispersal similar to *E. coli* Phollow virocells (Extended Data Fig. 6c). Following the colonization and treatment scheme in Fig. 4b, *Plesiomonas* also gave rise to clouds of virions throughout the gut (Fig. 4g and Supplementary Movie 4). However, a striking activity displayed by *Plesiomonas* Phollow phages is that they appeared to be quickly internalized by cells lining the gut (Supplementary Movie 5). We identified one of the intestinal cell types to be enteroendocrine cells (EECs), which we confirmed using *nkx2.2a*:*megfp* transgenic zebrafish^33,34^ (Fig. 4h and Supplementary Movie 6). We further observed that *Plesiomonas*-derived virions rapidly disseminate from the gut to extraintestinal sites. Most notably, virions accumulated in the liver within lipid droplet-like compartments (Fig. 4i). Moreover, patches of viral particles (and possibly fluorescent cell debris) were found in the vasculature, including within blood vessels of the brain (Fig. 4j). Finally, like *E. coli* virions, those from *Plesiomonas* turned over rapidly and were no longer detected within the gut or extraintestinal tissues 24h-post trimethoprim treatment. Together, these experiments, expedited by Phollow, show that P2-like phages from different bacterial hosts can share overlapping characteristics but that they also display phage or bacterial strain-dependent activities.

### Tracing interbacterial phage transmission in vitro

To monitor interbacterial phage transmission in vitro using Phollow, we first identified three fluorescent tags that enabled simultaneous visualization of intracellular and extracellular virions, and infection of new bacterial hosts (Fig. 5a–c). We then assembled a three-member community comprising mNG *E. coli* Phollow virocells and two *E. coli* target cells that are susceptible to P2 phage infection and that express mKate2 or mTFP SpyCatcher proteins (Fig. 5d). Treating this community with MMC induced lytic replication and dispersal of mNG Phollow phages, which could be seen landing on and replicating within each target cell type (Fig. 5e, insets i,ii). Strikingly, mKate2 and mTFP Phollow phages were also found landing on each cell type within the community, highlighting the potential for phage outbreaks to fuel onward transmission (Fig. 5e, insets iii–vi).

**Figure 5.**
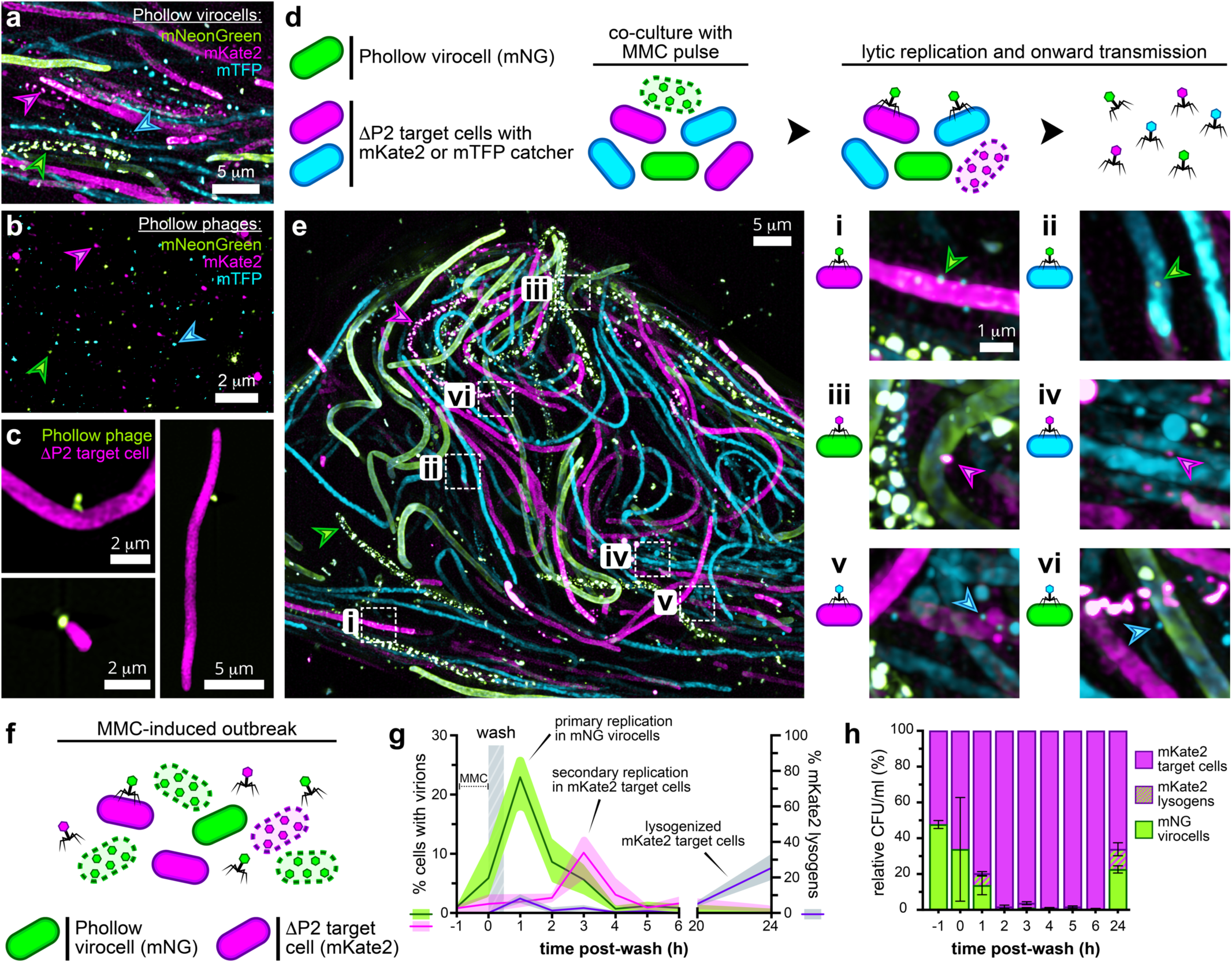
Tracing interbacterial phage transmission in vitro. **(a)** Maximum intensity projection image of a mixed culture containing mNeonGreen, mKate2, and mTFP *E. coli* Phollow virocells treated with MMC. Green (mNeonGreen), magenta (mKate2), and cyan (mTFP) arrowheads indicate cells harboring lytically replicating Phollow phage. **(b)** Maximum intensity projection image of purified mNeonGreen, mKate2, and mTFP Phollow phage derived from panel a. Green (mNeonGreen), magenta (mKate2), and cyan (mTFP) arrowheads indicate each of the Phollow phage types. **(c)** Three example maximum intensity projection images of mNeonGreen Phollow phage binding to an mKate2 *E. coli* target cell cured of its native P2 prophage (ΔP2). **(d)** Diagram of three-member community (left) and MMC induction scheme (right) for imaging interbacterial phage transmission. “MMC pulse” refers to a 1h MMC treatment followed by a wash step as described in Fig. 2 b,d. **(e)** Maximum intensity projection image of the three-member community diagrammed in panel d following MMC-treatment. Green arrowhead indicates an mNG Phollow virocell harboring lytically replicating phage. Magenta arrowhead indicates an mKate2 target cell harboring lytically replicating phage, which indicates an instance of interbacterial transmission. (right) Insets show mNG (i,ii), mKate2 (iii,iv), and mTFP (v,vi) Phollow phage landing on each bacterial community member. **(f)** Diagram of two-member community for tracing interbacterial phage transmission dynamics. **(g)** Left y-axis: Quantification of lytic replication in mNG virocells (green line) and mKate2 target cells (magenta line) by imaging flow cytometry. Right y-axis: Enumeration of lysogenized mKate2 target cells (purple line) as a percentage of total mKate2 target cells determined by differential plating. Shaded regions indicate ranges from 3 biological replicates performed at each time point. **(h)** Enumeration of community composition from panel g by differential plating. Data are presented as average relative abundances; bars indicate standard error of the mean (SEM) from across 3 biological replicates at each time point.

We next determined the capacity of Phollow to enable the quantitative tracking of phage replication and transmission. We induced an outbreak of Phollow phages in a two-member community of mNG *E. coli* Phollow virocells and mKate2 *E. coli* target cells (Fig. 5f). We then followed viral replication and transmission over time, along with the abundance of each community member using imaging flow cytometry and plating (Fig. 5g,h). MMC induced an acute wave of mNG phage lytic replication and dispersal that coincided with a sharp decline in the mNG virocell population (Fig. 5g,h). mNG phages infecting mKate2 target cells triggered a subsequent wave of mKate2 phage lytic replication, with lytic replication subsiding in both populations by 24h (Fig. 5g). Intriguingly, at 24h the mNG virocell population rebounded to 22.6% of the community whereas mKate2 target cells made up 66.1%. Moreover, a new population had emerged: an mKate2 virocell generated by lysogeny, which made up 25.4% of all mKate2 cells and 11.2% of the total community (Fig. 5g,h). Ultimately, this experiment shows how Phollow can reveal the underlying transmission dynamics that shape bacterial communities.

### Mapping spatiotemporal dynamics of phage replication regimes within the gut

In the vertebrate gut, resident communities are spatially structured and dynamic but the relationship between community organization and phage replication regimes is largely undefined. We therefore used Phollow to track interbacterial phage transmission during an outbreak of lytic replication within the zebrafish gut. We colonized four-day old germ-free zebrafish with a two-member bacterial community composed of AausFP1 *E. coli* Phollow virocells and mKate2 *E. coli* target cells (Fig. 6a). We then monitored community and phage replication dynamics in response to trimethoprim by live imaging and plating of homogenized intestinal tissues. Unexpectedly, prior to treatment, AausFP1 virocells displayed a colonization advantage over mKate2 target cells (Fig. 6b, “untreated”), with both populations showing strong spatial segregation (Fig. 6c, top). Lytic replication in AausFP1 virocells could be detected as early as 2h post-treatment and was followed by a rapid reconfiguration of community organization (Fig. 6c, middle). By 4h post-treatment, the two-member community displayed a striking level of spatial mixing (Fig. 6c, bottom). Surprisingly, the relative abundances of each member at the 4h time point were mostly unchanged, and we did not detect instances of lysogeny in mKate2 target cells (Fig. 6b, “Trimethoprim’’). In addition, only AausFP1 virions had been expelled into the water (Fig. 6d). These results suggest that immediately after trimethoprim treatment, there was likely little to no interbacterial phage transmission within the gut.

**Figure 6.**
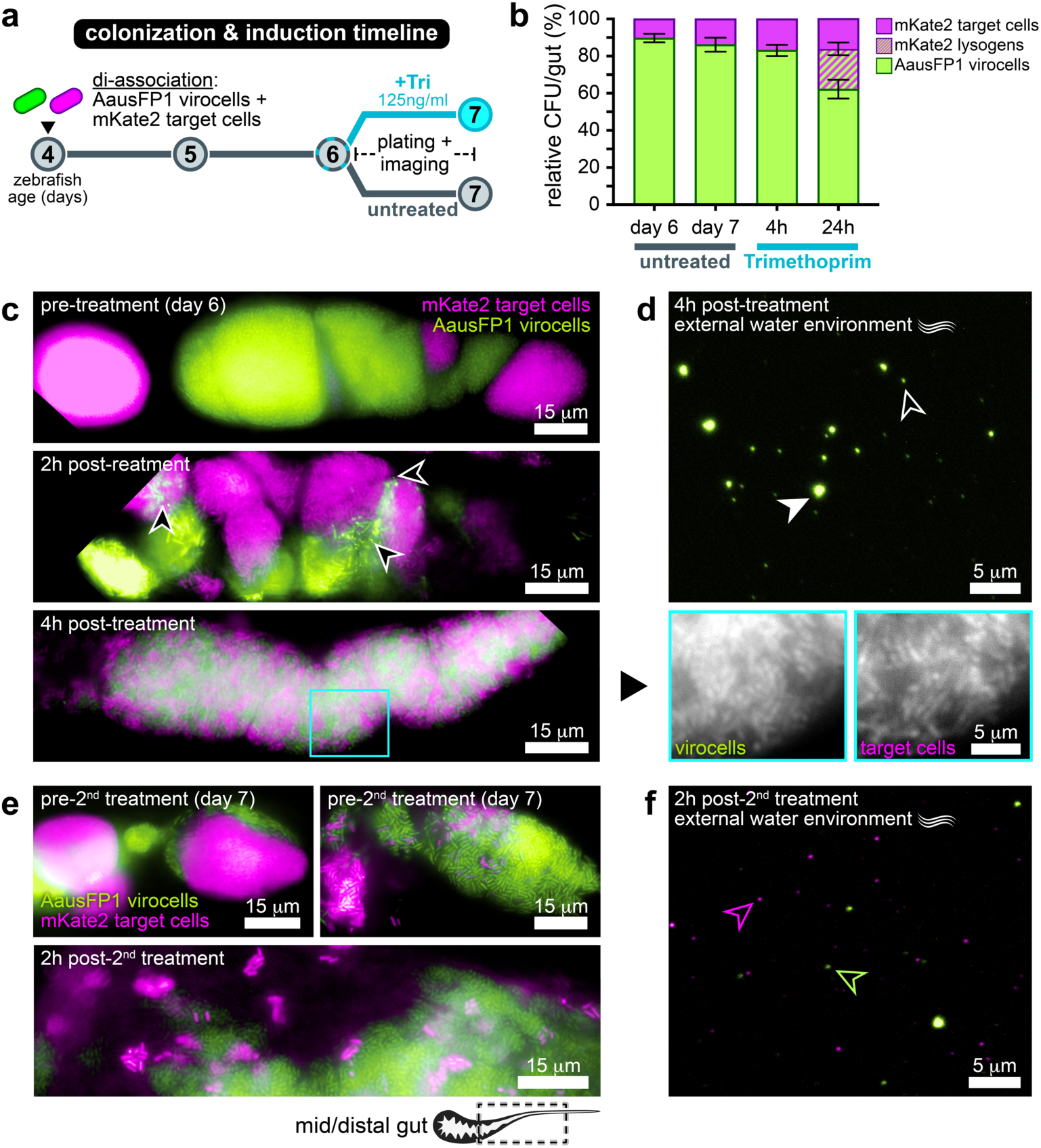
Mapping spatiotemporal dynamics of phage replication regimes within the gut. **(a)** Diagram of bacterial colonization and antibiotic induction timeline. “Tri” = Trimethoprim. **(b)** Enumeration of community composition by differential plating. Data are presented as average relative abundances; bars indicate standard error of the mean (SEM) from across 3 biological replicates at each time point and condition. **(c)** Fluorescent microscopy images of AausFP1 virocell/mKate2 target cell gut communities prior to (top), 2h post- (middle), and 4h post-trimethoprim treatment. Each image was taken from different zebrafish hosts. Black arrowheads in the middle image indicate areas containing lytically replicating phage. Inset in the bottom panel shows the AausFP1 virocell and mKate2 target cell channels separately to highlight degree of community mixing. **(d)** Representative fluorescence microscopy image of a water sample at 4h post-trimethoprim treatment. White arrowhead indicates a virion aggregate; Black arrowhead indicates a single viral particle. **(e)** Fluorescent microscopy images of AausFP1 virocell/mKate2 target cell gut communities prior to (top) and 2h after (bottom) a 2^nd^ post-trimethoprim treatment. Each image is from different zebrafish hosts. **(f)** Representative fluorescence microscopy image of a water sample 2h after a 2^nd^ trimethoprim treatment. Magenta arrowhead indicates an mKate2 Phollow phage virion; green arrowhead indicates an AausFP1 Phollow phage virion.

We considered the possibility that phage transmission in the gut was delayed compared to the dynamics we observed in vitro (Fig. 5g,h). No viral particles were found within the gut 24h post-treatment, but intriguingly, communities showed substantial recovery of a pre-treatment spatial structure that was marked by strong segregation of each bacterial population (Fig. 6e, top). Probing community composition revealed the presence of mKate2 virocells (formed by lysogeny) with a slight decline in AausFP1 virocell abundance (Fig. 6b, “Trimethoprim”). This observation indicates that between 4h and 24h, horizontal transmission of AausFP1 Phollow phages had occurred. In contrast, in the absence of trimethoprim, we did not detect lysogenized mKate2 virocells (Fig. 6b, “untreated”), which emphasizes the role of the drug-induced phage outbreak in sparking interbacterial spread. To further confirm that horizontal transmission had occurred, we induced a second wave of lytic replication by treating animals with another round of trimethoprim. We again observed immediate spatial fragmentation of the bacterial community (Fig. 6e, bottom), but during this second treatment, both AausFP1 and mKate2 Phollow phages were expelled into the water (Fig. 6f). We surmise that the source of mKate2 virions was likely mKate2 virocells formed during the first wave of AausFP1 phage lytic replication. Altogether, our experiments begin to reveal that there exists a co-dynamic between the spatial organization of gut bacterial communities and phage dispersal and transmission.

## DISCUSSION

Previous approaches using fluorescently marked virions have uncovered amazing details of phage biology^24,35–38^. However, comprehending the impacts of phages on microbiomes requires additional strategies for capturing phage replication and transmission across space, time, and complexity. Our solution is Phollow, which blends new genetic tools and experimental models to explore phage biology in unique ways. Combinatorial in vivo tagging using the SpyTag:SpyCatcher system makes it possible to investigate phage replication as well as dispersal and interbacterial transmission (Fig. 1). We showed that Phollow phages are amenable to both live imaging and flow virometry techniques (Fig. 2,3), which has the potential to facilitate novel experimental schemes aimed at dissecting the epidemiology of phage outbreaks (for example, as in Fig. 5). Emphasizing this potential, Phollow exposed new aspects of P2 phage replication; namely, the formation of virion aggregates that rapidly disassemble into single particles upon bacterial lysis (Supplementary Movie 2). The function of P2 virion aggregates is unknown but we posit that they may have a role in the evasion of antiphage defense systems similar to the nucleus-like structures formed by jumbo phages^39^. It is also possible that aggregates act to shield virions and preserve infectivity when exposed to harsh extracellular environments^40,41^.

Visualizing lytic replication within the gut revealed previously unseen patterns of dispersal and biogeography (Fig. 4). Antibiotic-induced outbreaks occurred on rapid timescales and generated clouds of virions that often filled the luminal space (Supplementary Movie 4). However, the residency of virions within the gut was surprisingly short-lived. We surmise that the fast turnover of virions is driven by surges of peristaltic activity, which are known to purge bacterial populations and control community composition^26,42^. Equally surprising was the speed with which phage outbreaks sparked transformations in the spatial structure of gut bacterial communities (Fig. 6c). It is likely that spatial structure within a community regulates horizontal transmission of phages, but additional work is needed to fully understand this relationship. However, our observations highlight the intriguing possibility that phage replication regimes may create patterns of organization by affecting bacterial growth and death across a community. Future studies will use Phollow to decipher the interplay between community organization and phage outbreaks and probe the impact on interactions across a microbiome.

Our finding that Phollow phages from *Plesiomonas* readily associate with and disseminate across zebrafish tissues provides new opportunities for studying how prokaryotic viruses interact with and modulate eukaryotic biology^43–46^ (Fig. 4h–j). In mammals, translocation of phages across the gut epithelium is suggested to be an active and ongoing process^47^. Here we find that EECs and perhaps other cell types may be a significant conduit for phage dissemination (Supplementary Movies 5 and 6). The prospect that EECs are involved in phage translocation is further compelling given that they directly sense luminal nutrients and microbial products, and respond by modulating metabolic and immunological signaling throughout the body^48^. Beyond the gut, the presence of virions within the zebrafish liver (Fig. 4i) is consistent with studies showing that phages injected into the bloodstream are filtered by hepatocytes^49,50^. Moreover, in pigs and macaques, the DNA of resident gut phages can be recovered from parenchymal organs, including the liver^51^. Notably, it was recently shown that some phages have immunomodulatory and proinflammatory potential^43,46,52,53^. Therefore, it will be interesting to investigate whether *Plesiomonas*-derived phages also influence inflammatory tone.

Ultimately, the strength of Phollow is that it enables the nanoscopic lives of bacterial viruses to be directly visualized. Although we used Phollow to illuminate facets of P2 phage biology, we expect that it can also accelerate the exploration of other phage lineages. In particular, the immense diversity of phages combined with the dizzying amount of viral genetic dark matter remaining to be functionally annotated makes it nearly impossible to predict the activity of a given phage within its natural environment. We think that Phollow can help overcome this challenge. We note, however, that Phollow does have some limitations. As with any approach involving genetic modifications, there is a chance that fluorescently marking virions with Phollow disrupts their native functions. For example, phages that rely on capsid-mediated interactions may be especially sensitive to Phollow tagging procedures^54,55^. However, this could be circumvented using sparse labeling strategies that preserve capsid functionality. In addition, live imaging techniques alone are limited in their ability to extract information from biological systems. Therefore, we envision that it will require a combination of conventional and novel approaches, as well as new model systems, to piece together the dynamic and nested life cycles of phages within the context of complex microbiomes.

## ACKNOWLEDGMENTS

Research reported in this publication was supported by the National Institute of Allergy and Infectious Diseases of the National Institutes of Health under award number DP2AI154420 to T.J.W. The content is solely the responsibility of the authors and does not necessarily represent the official views of the National Institutes of Health. The Gonen lab is supported by The National Institute of General Medical Sciences, grant R35-GM142797. N.U. and N.H.R. were supported by NIH supplements (R00 HD098330 04S1 to N.U. and DP2AI154420-03S1 to N.H.R). D. S. S. and C.T. were supported by a graduate fellowship from the Graduate Assistance in Areas of National Need (GAANN) (P200A210024) provided by the U.S. Department of Education. M.Z. was supported with a PostDoc.Mobility fellowship (P500PB_214357) from the Swiss National Science Foundation.

This study was made possible in part through access to the Optical Biology Core Facility of the Developmental Biology Center, a shared resource supported by the Cancer Center Support Grant (CA-62203), the National Institutes of Health grant (NIH S10OD032327-01) and the Center for Complex Biological Systems Support Grant (GM-076516) at the University of California, Irvine.

We thank the UC Irvine Center for Virus Research for Spectral Flow Cytometry Access and the support of the Chao Family Comprehensive Cancer Center Flow Cytometry shared facility, supported by the National Cancer Institute of the National Institutes of Health under award number P30CA062203.

pDEST14-SpyCatcher (Addgene plasmid # 35044), SpyCatcher003-sfGFP (Addgene plasmid # 133449) and pET28a-SnoopCatcher (Addgene plasmid # 72322) were a gift from Mark Howarth; TOPO-mKate2 (Addgene plasmid # 68441) was a gift from Tyler Jacks; pNCST-AausFP1 (Addgene plasmid # 129499) was a gift from Nathan Shaner; mCitrine-pBAD (Addgene plasmid # 54723) was a gift from Robert Campbell & Michael Davidson & Oliver Griesbeck & Roger Tsien; mScarlet_C1 (Addgene plasmid # 85042) was a gift from Dorus Gadella; mTFP1-pBAD (Addgene plasmid # 54553) was a gift from Robert Campbell & Michael Davidson.

Finally, we are grateful to Dr. Michael Parsons (University of California, Irvine) for his incredible generosity and support with our zebrafish colony since the beginning of the COVID-19 pandemic in 2020. We also thank Dr. Parsons for sharing the TgBAC(*nkx2.2a*:*megfp*) zebrafish line.

## AUTHOR CONTRIBUTIONS

*Conceptualization*: L.O., E.T.W, and T.J.W

*Formal analysis*: L.O., E.T.W, M.Z., and T.J.W

*Funding acquisition*: T.J.W

*Investigation*: L.O., E.T.W, M.Z., and T.J.W

*Methodology*: L.O., E.T.W, M.Z., M.B., N.H.R., D.S.S., N.U., J.A.A., C.T., C.K., and T.J.W

*Supervision*: S.G., D.N., and T.J.W

*Visualization*: L.O. and T.J.W

*Writing – original draft*: L.O., and T.J.W

*Writing – review & editing*: All authors

## SUPPLEMENTARY MOVIE LEGENDS

**Supplementary Movie 1. Examples of intracellular Phollow phage organization.** 360° view of images presented in Fig. 2e and Extended Data Fig. 3 highlighting patterns of intracellular virion organization. Virions show two prevalent patterns of organization: a “scattered” distribution and a “serpentine” distribution. Images are pseudo-colored according to z-depth, representing a total of 1.36 μm.

**Supplementary Movie 2. Time-lapse movie depicting Phollow phage induction, assembly, and dispersal.** Time-series images of *E. coli* HS Phollow virocells growing on agar pads containing MMC. Optical frames were generated from four tile-scanned and merged fields of view, acquired at regular 10-minute intervals over a period of 370 minutes. The time series starts at 0 minutes, where individual bacterial cells appear as small white rods. As time progresses, these cells undergo filamentation, a hallmark morphological change indicative of the SOS response induced by MMC genotoxicity. Subsequently, a subpopulation of bacterial cells displays the formation of intracellular fluorescent viral foci followed by explosive cell lysis. Upon lysis, Phollow phage virions can be seen dispersing throughout the extracellular milieu.

Supplementary Movie 3. Three-dimensional scan through the body of a 7-day old larval zebrafish colonized by *E. coli* HS Phollow virocells. Optical frames were generated from 297 tile-scanned and merged images acquired by super-resolution microscopy. DNA (magenta, JF549 dye) and actin (purple, CF405-phalloidin) highlight zebrafish anatomy and tissues, bacteria (green) localize as aggregates in the midgut.

**Supplementary Movie 4. Spatial distribution of Phollow phage virions within the zebrafish gut.** Three-dimensional scan through the anterior region of a 6-day old larval zebrafish intestine colonized by AausFP1 *Plesiomonas* Phollow virocells. The video indicates the presence of bacteria and viral clouds throughout the luminal space. Images were acquired 4h post-trimethoprim treatment, during an outbreak of phage lytic replication. DNA (magenta, JF549 dye) and actin (purple, CF405-phalloidin) highlight the intestinal structure.

**Supplementary Movie 5. Phollow phage virions are dynamic and closely associate with the intestinal mucosa.** Live imaging of the anterior region of a 6-day old larval zebrafish intestine colonized by AausFP1 *Plesiomonas* Phollow virocells. The video shows the close association with and possible translocation of virions across the gut epithelium.

**Supplementary Movie 6. Example of internalization of Phollow phage virions by enteroendocrine cells.** Three-dimensional reconstruction of intestinal tissue highlighting the co-localization of viral particles (grayscale) within an enteroendocrine cell (EEC, green). The membranes of EECs are marked using TgBAC(*nkx2.2a*:*megfp*) reporter fish and DNA is stained with Hoechst 33342 dye (magenta). Surface rendering and 360° view were generated from z-stack images using Imaris.

---

**Extended Data Fig. 1.**
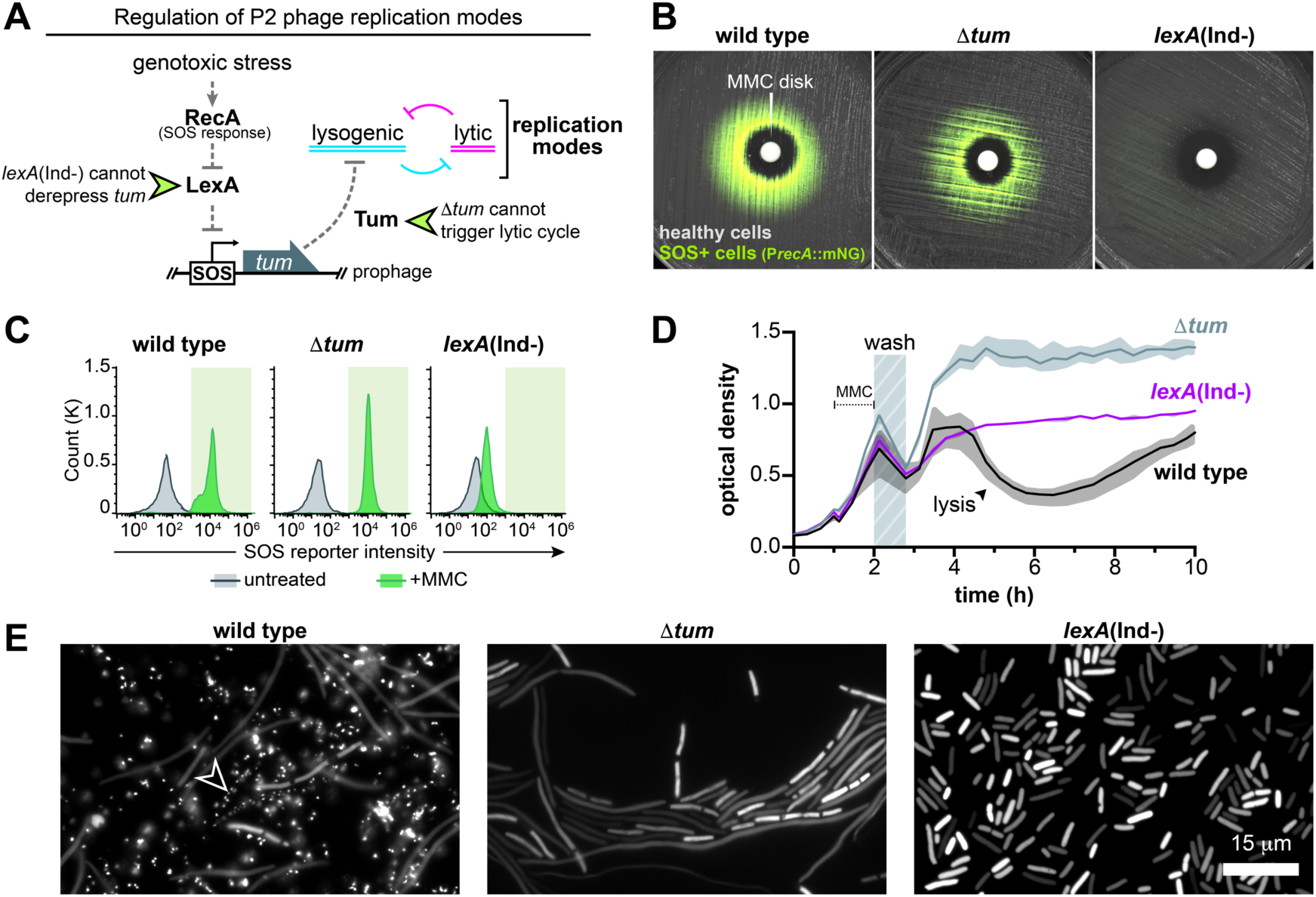
P2 phage replication modes in *E. coli* HS are controlled by the DNA damage “SOS” response. **(a)** Diagram of the signaling pathway controlling P2 phage replication modes in response to genotoxic stress. DNA damage activates RecA, which mediates the cleavage of the LexA repressor protein. LexA controls the expression of genes within the SOS regulon by binding to operator sequences known as SOS boxes. Within the P2 prophage genome, the LexA-controlled *tum* gene becomes derepressed leading to expression of the Tum antirepressor protein. Tum toggles lytic replication by inhibiting pro-lysogenic regulators encoded by the prophage. Bacterial mutants encoding a non-cleavable *lexA* allele (*lexA*(Ind-)) are unable to derepress *tum* and thus, are unable to trigger lytic replication. Likewise, prophage mutants lacking *tum* (Δ*tum*), are also unable to trigger lytic replication. **(b)** Disk diffusion tests visualizing activation of the SOS response by MMC. A fluorescent SOS reporter was constructed by fusing the promoter of the *recA* gene (P*recA*) to an open reading frame encoding mNeonGreen (mNG). *E. coli* HS strains carrying the SOS reporter were spread onto agar plates at a density sufficient to produce a lawn of growth. Antibiotic assay disks were then placed in the center of the agar plate and loaded with MMC. Plates were incubated overnight at 37°C and imaged using a fluorescence stereomicroscope. MMC generated a zone of inhibition around the assay disk for each strain. However, only the wild-type and Δ*tum* strains displayed SOS reporter activity in cells adjacent to the zone of inhibition. As expected, the *lexA*(Ind-) mutant strain did not display SOS reporter activation. **(c)** Quantification of SOS reporter activity following a MMC pulse via flow cytometry. Bacterial cells carrying the fluorescent SOS reporter were analyzed 1h after washing MMC from the cell cultures. Histograms display the distribution of SOS reporter intensity in untreated (gray) and MMC-treated (green) cultures, with the green shaded area indicating strong reporter activity. The MMC pulse effectively induced SOS reporter activity in wild-type and Δ*tum* strains, whereas the *lexA*(Ind-) mutant exhibited a minor shift in intensity. **(d)** Representative induction (or “lysis”) curve of P2 phage lytic replication in *E. coli* HS. The optical density of cultures (at 600 nm) containing either wild-type (black line), Δ*tum* (gray line) or *lexA*(Ind-) (purple line) strains was monitored prior to and following a 1h MMC treatment. Shaded regions represent the range of optical density measurements from three technical replicates. MMC was washed from cultures at 2h (hashed bar), which prevents interference of phage lytic replication. Washing also produces a slight decrease in cell density due to some cell loss. An abrupt drop in optical density of the wild-type culture at 4h marks widespread lysis, which is not observed in Δtum or *lexA*(Ind-) cultures, suggesting attenuation of lytic replication in these strains. **(e)** Fluorescence microscopy images of wild-type, Δ*tum*, and *lexA*(Ind-) Phollow virocell strains 1h post-MMC treatment. Wild-type cells show extensive filamentation and formation of viral foci (arrowhead), which are not displayed by Δ*tum* and *lexA*(Ind-) mutant strains. Additionally, *lexA*(Ind-) cells do not exhibit cell filamentation due to an impaired SOS response. These results demonstrate the necessity of both an active SOS signaling pathway and *tum* expression to toggle P2 phage lytic replication.

**Extended Data Fig. 2.**
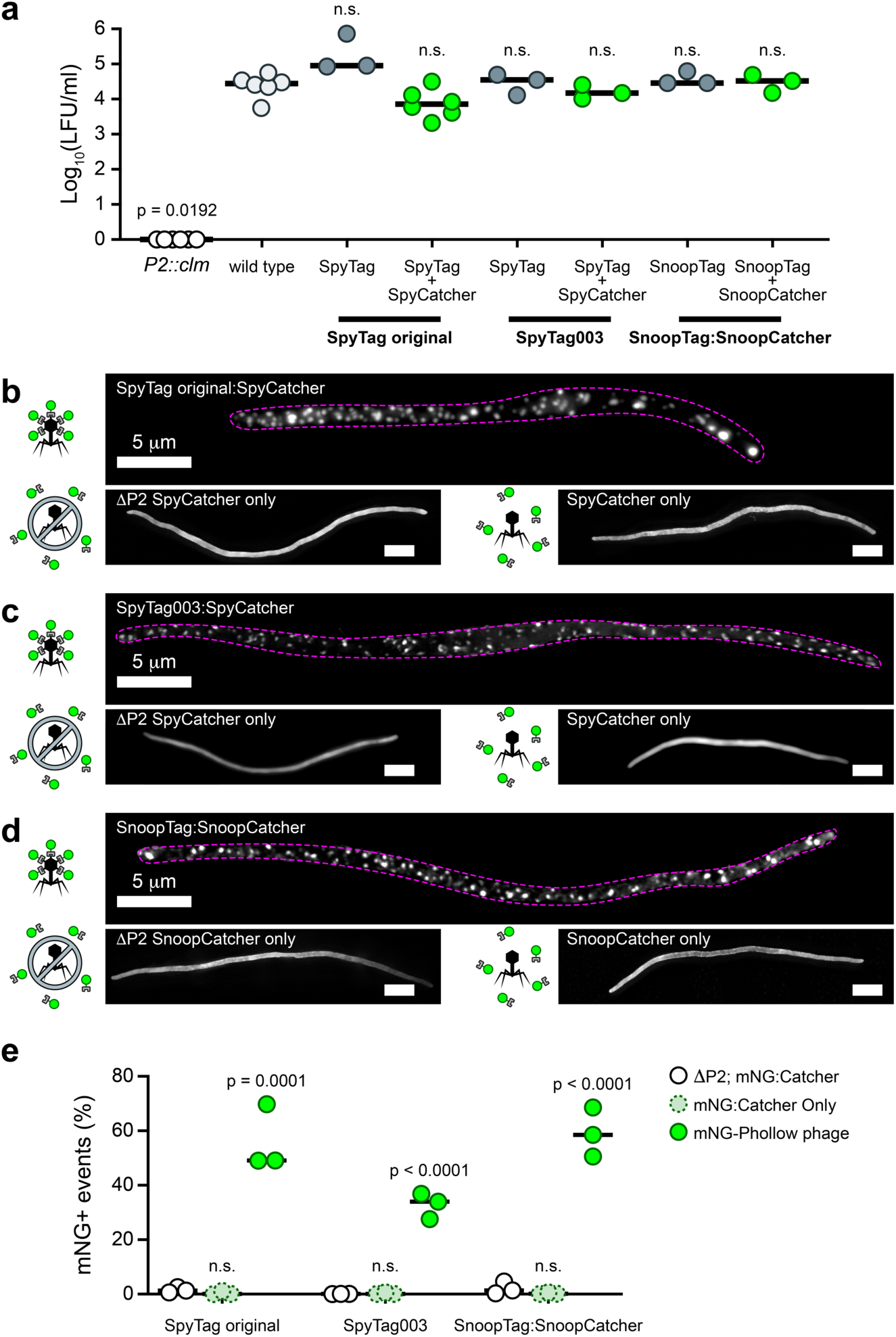
Characterization of Phollow phage tagging variants. **(a)** The infectivity of wild-type and different Phollow phage variants was assessed by measuring lysogen-forming units (LFU) generated from infection of an *E. coli* HS target cell cured of its resident P2 prophage. Lysogeny was monitored using prophages carrying a chloramphenicol resistance gene. *P2::clm* is an *E. coli* HS strain that carries a chloramphenicol resistance marker in place of the P2 prophage genome and is used to control for phage-independent transfer of chloramphenicol resistance. Bars denote medians and each circle represents an independent biological replicate. Significant differences compared to “wild type” were determined by one-way ANOVA using a Kruskal-Wallis and Dunn’s multiple comparisons test (p < 0.05; n.s. = not significant). **(b-d)** Comparison of viral foci production and organization across Phollow phage variants: (b) SpyTag original:SpyCatcher; (c) SpyTag0003:SpyCatcher; (d) SnoopTag:SnoopCatcher. The top of each panel shows a representative maximum intensity projection image of viral foci organization within Phollow virocells capable of generating fully tagged Phollow phages (a dashed magenta line highlights the cell perimeter). The bottom left of each panel shows a representative maximum intensity projection image of Spy- or SnoopCatcher-based fluorescence in the absence of a P2 prophage (ΔP2). The bottom right of each panel shows a representative maximum intensity projection image of Spy- or SnoopCatcher-based fluorescence in the presence of a lytically replicating P2 phage that lacks a Spy- or SnoopTag, which is thus unlabeled. No viral foci form in the absence of a fully tagged Phollow phage. **(e)** Plot shows the functionality of each Phollow phage variant to be used for flow virometry. Data are presented as the percentage of mNeonGreen (mNG) positive events. Statistical differences compared to the “ΔP2; mNG:catcher” control strain within each variant group were determined by an ordinary one-way ANOVA using a Dunn’s multiple comparisons test (p < 0.05; n.s. = not significant).

**Extended Data Fig. 3.**
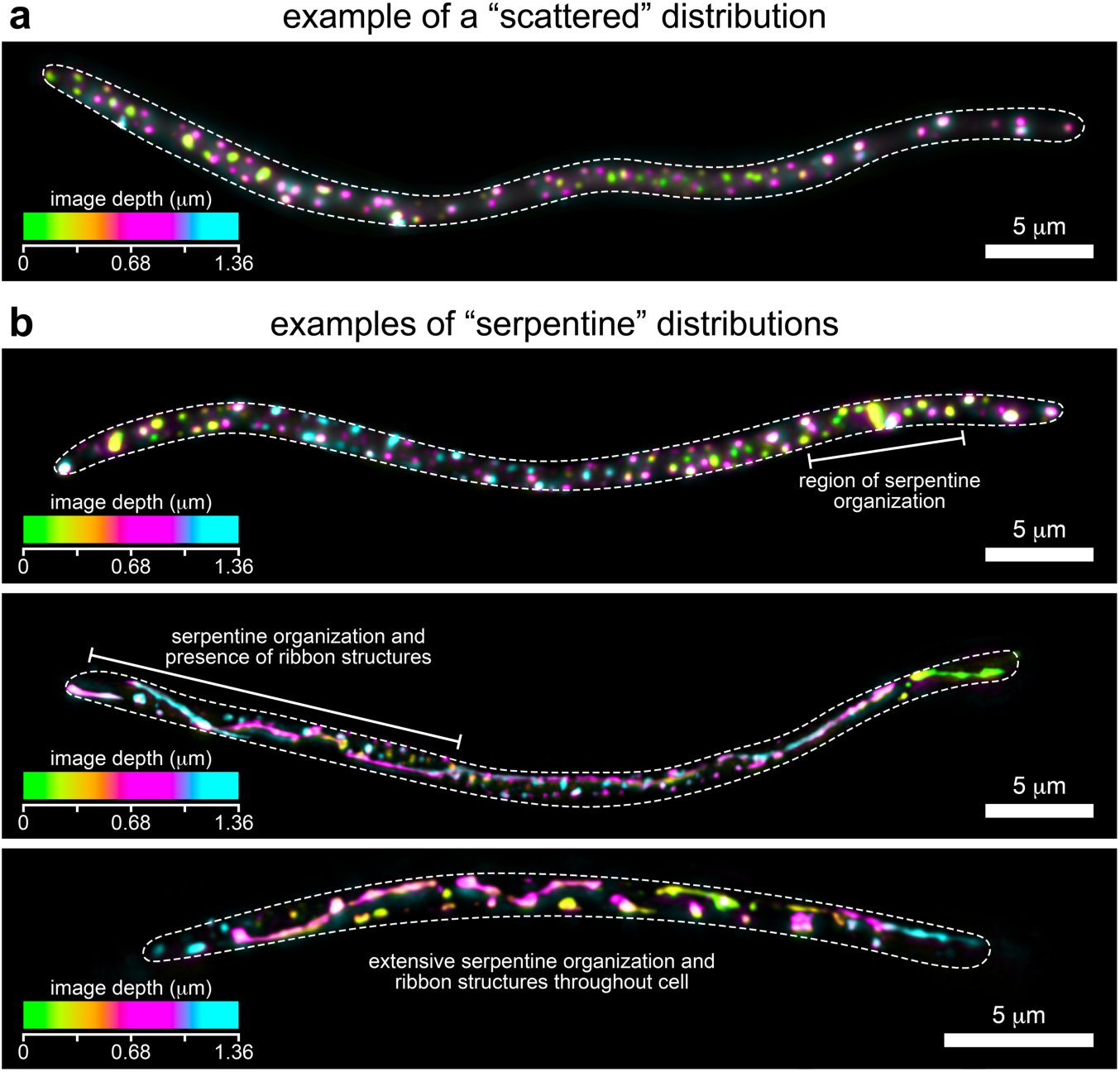
Examples of intracellular Phollow phage organization. **(a-b)** Z-projected images of bacterial cells harboring viral foci. Images highlight two patterns of virion organization throughout the intracellular space: (a) a scattered distribution and (b) a serpentine pattern with the presence of ribbon structures. Images are pseudo-colored according to z-depth, representing a total of 1.36 μm. A white dashed line marks the cell perimeter.

**Extended Data Fig. 4.**
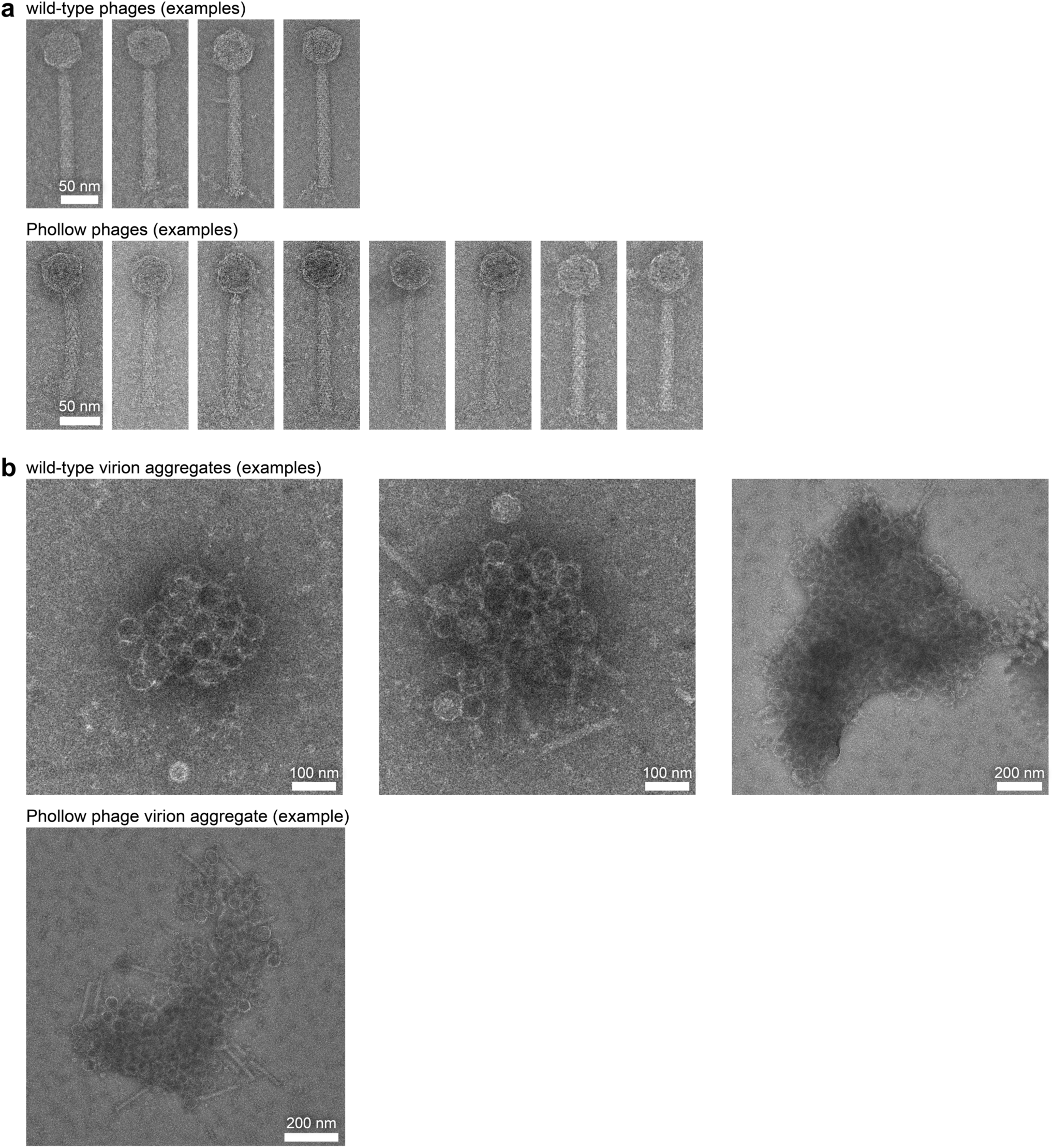
Transmission electron microscopy (TEM) of negatively stained P2 phage virions and virion aggregates. **(a)** Representative TEM micrographs of unmodified wild-type (top) and mNeonGreen Phollow phage virions (bottom). **(b)** TEM micrographs of extracellular virion aggregates from unmodified wild-type P2 phage (top) and mNeonGreen Phollow phage (bottom).

**Extended Data Fig. 5.**
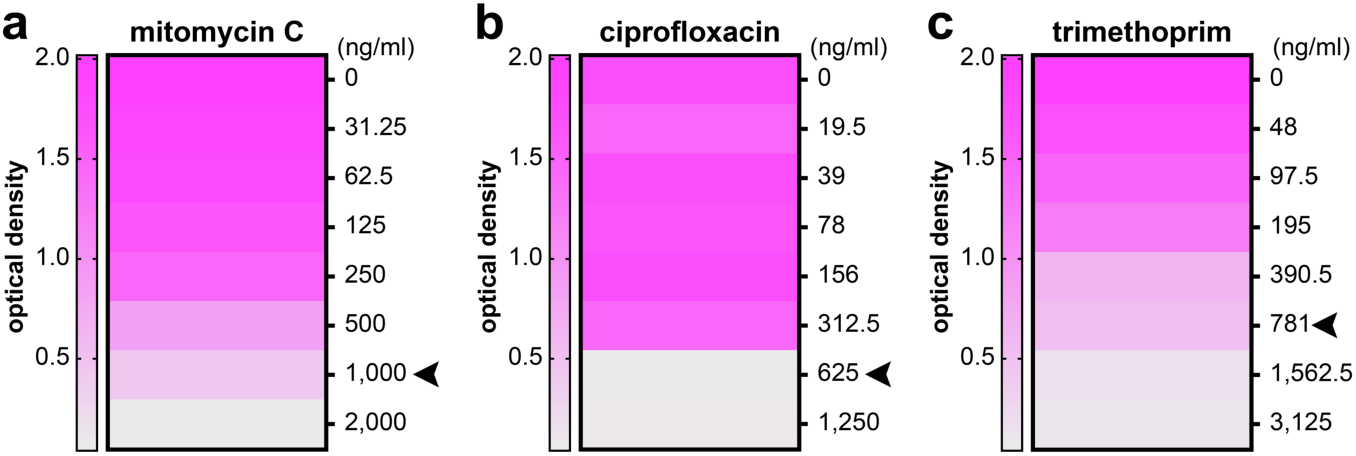
Determination of minimum inhibitory concentrations of genotoxic antibiotics for *E. coli* HS. **(a-c)** Optical density at 600 nm measurements of *E. coli* HS cultures in response to various concentrations of (a) MMC, (b) ciprofloxacin, and (c) trimethoprim. Data represent the average of three independent endpoint measurements after 24 hours of incubation at 37°C, plotted as heatmaps. Arrowheads in each panel indicate the minimum inhibitory concentration (MIC), defined as the minimum concentration required to significantly impair bacterial growth without complete eradication.

**Extended Data Fig. 6.**
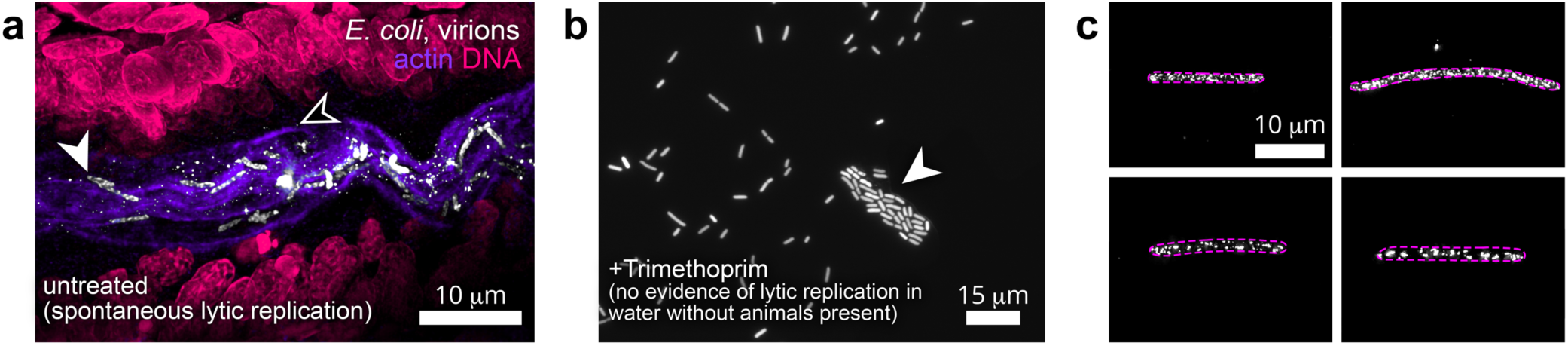
Characterization of phage outbreaks within the vertebrate gut. **(a)** Maximum intensity projection image showing viral particles derived from untreated *E. coli* HS Phollow virocells undergoing spontaneous phage lytic replication within the intestine. White arrowhead marks bacterial cells; black arrowhead marks viral particles. **(b)** Representative fluorescent microscopy images of AausFP1 *E. coli* HS Phollow virocell in zebrafish embryo media (EM) in the absence of fish. Bacteria were separated from fish immediately before the experiment to allow for the diffusion of fish-derived nutrients and other factors. Subsequently, bacteria were treated with trimethoprim and incubated at 28.5°C for 4h. Bacterial cells under these conditions do not display viral foci or signs of phage lytic replication (white arrowhead). **(c)** Maximum intensity projection images of MMC-induced *Plesiomonas* Phollow virocells. Each panel shows a representative image of fluorescent viral foci production and organization, which are similar to those in *E. coli* Phollow virocells. Magenta dashed lines mark the cell perimeter.

## MATERIALS AND METHODS

### Animal care

All experiments with zebrafish were done in accordance with protocols approved by the University of California, Irvine Institutional Animal Care and Use Committee (protocol #AUP-23-126) and followed standard protocols^56^. Specific handling and housing of animals during experiments are described in detail under the section “Gnotobiology”. All zebrafish used in this study were larvae, between the ages of 4- and 7-days post fertilization. Sex differentiation occurs later in zebrafish development and thus was not a factor in our experiments. Zebrafish lines used in this study included wild-type AB and zebrafish carrying the TgBAC(*nkx2.2a*:*megfp*) transgene which expresses a membrane-tethered enhanced green fluorescent protein (mEGFP) under control of *nkx2.2a* regulatory sequences carried on a bacterial artificial chromosome (BAC)^33,34^.

### Gnotobiology

Wild-type (AB) and transgenic TgBAC(*nkx2.2a*:*megfp*)^33,34^ zebrafish embryos were derived germ-free (GF) and colonized with bacterial strains as previously described^57^ with slight modifications. Briefly, fertilized eggs from adult mating pairs were harvested and incubated in sterile embryo media (EM) containing ampicillin (100 μg/ml), gentamicin (10 μg/mL), amphotericin B (250 ng/mL), tetracycline (1 μg/mL), and chloramphenicol (1 μg/mL) for ∼6h. Embryos were then washed in EM containing 0.1% polyvinylpyrrolidone-iodine followed by EM containing 0.003% sodium hypochlorite. Sterilized embryos were distributed into T25 tissue culture flasks containing 15 ml sterile EM at a density of one embryo per milliliter and incubated at 28.5°C prior to bacterial colonization. Embryos were sustained on yolk-derived nutrients and were not fed during experiments. For bacterial association studies, bacterial strains were grown overnight as described in “Bacterial strains and culture”. Bacteria were then prepared for inoculation by pelleting 1 mL of culture for 2 min at 7,000 × g and washing once in sterile EM. Bacterial strains were individually added to the water of single flasks containing 4-day-old larval zebrafish at a final density of 10^6^ bacteria/mL.

### Bacterial strains and culture

All wild and recombinant bacterial strains used or created in this study are listed in Extended Data Table 1 and Table 3. Archived stocks of bacteria were maintained in 25% glycerol at −80°C. Prior to manipulations or experiments, bacteria were directly inoculated into 5 mL TSB media (TSB; MP Biomedicals, VWR) and grown for ∼16 h (overnight) with shaking at 37°C, except for *Plesiomonas* ZOR0011, which was grown at 30°C. For growth on solid media, tryptic soy agar (TSA; Hardy Diagnostics, VWR) was used. When specified, media was supplemented with 10 µg/mL gentamicin or 20 µg/mL chloramphenicol.

### Molecular techniques and strain construction

Plasmids were constructed using standard molecular cloning techniques as previously described^30^ and in vivo assembly (IVA)^58^, using primers listed in Extended Data Table 4. All engineering plasmids and recombinant strains were sequence- and/or PCR-confirmed.

#### Phollow phage and virocell engineering

SpyCatcher^17^ (Addgene 35044), SpyCatcher003^21^ (Addgene 133449), and SnoopCatcher^22^ (Addgene 72322) plasmids were purchased from Addgene and IVA was used to fuse each Catcher gene to the N-terminus of individual fluorescent proteins, along with a GSSGS linker. Catcher fusions were cloned into the pXS^30^ expression scaffold. Next, Catcher fusion genes along with the constitutive pTac promoter and transcriptional terminator contained within pXS were subcloned into pTn7xTS^30^ for transposon-based insertion into the *attTn7* site of the bacterial chromosome. SpyTag, SpyTag003, and SnoopTag were fused to the C-terminus of the P2 phage major capsid protein (GpN) with a SGGGSG linker in pAX1^30^ using IVA. The resulting plasmids were used for allelic exchange to generate a markerless modification of the gene encoding GpN within the prophage genome in the chromosome of *E. coli* HS or *Plesiomonas* ZOR0011 as previously described^30^. Further details on plasmid design and construction are provided in Supplementary File 1.

#### Creation of ΔP2 “target cell”, Δtum, and lexA(Ind-) strains of E. coli HS

Mutant strains were created by markerless allelic exchange using pAX1^30^. For ΔP2 “target cell” strains used as recipients for P2 phage transmission experiments and infectivity (LFU) assays, allelic exchange cassettes were designed to delete the entire P2 prophage genome while reconstituting the attachment site (*attB*) on the bacterial chromosome to enable P2 phage integration and lysogeny. For the Δ*tum* mutant strain, an in-frame deletion of the prophage gene encoding the Tum antirepressor was made using an allelic exchange cassette that fused the *tum* start and stop codons. For the *lexA*(Ind-) strain, the *lexA* open reading frame was engineered using allelic exchange to encode a G85D residue mutation in the LexA protein (via a GGT to GAC codon change). See Supplementary File 1 for plasmid construction details.

#### Creation of fluorescent “SOS” reporter

A fluorescent SOS reporter was engineered by fusing the promoter region upstream of the *E. coli* HS *recA* gene (locus tag ECHS_RS13985) (P*recA*) with an open reading frame encoding mNeonGreen within a pTn7xTS vector. See Supplementary File 1 for plasmid construction details. The reporter was inserted into the *attTn7* site of the bacterial chromosome via Tn7 tagging as previously described^30^.

#### Creation of marked P2 phage variants for measuring infectivity by LFU assays

To measure the infectivity of *E. coli* HS P2 phage variants, a gene encoding resistance to chloramphenicol (from pKD3^59^) was inserted into the second moron region^19^ of the P2 prophage (between tail genes GpH and GpFI) using the lambda Red-mediated linear transformation system^59,60^. Briefly, the chloramphenicol resistance gene from the pKD3 plasmid was PCR-amplified with 40-base pair overhangs specific to the 5’ and 3’ ends of the second moron region. The resulting PCR amplicon was then introduced via electroporation into *E. coli* HS strains carrying pKM208, which encodes an IPTG (isopropyl-b-D-thiogalactopyranoside)-inducible lambda Red recombinase. Successful recombinants were selected on plates containing chloramphenicol (creating HS *P2 moron::clm*). As a control for LFU infectivity assays, an *E. coli* HS “donor” strain (HS *P2::clm*) was created by lambda Red recombination in which the chloramphenicol resistance gene was used to replace the entire P2 prophage.

### In vitro characterization of P2 phage induction

#### Induction of P2 phage lytic replication

Induction of the P2 phage was achieved by treating *E. coli* HS strains with a pulse of mitomycin C (MMC; Goldbio). Initially, overnight bacterial cultures were diluted 1:100 in fresh TSB media and incubated at 37°C with shaking for one hour. MMC was then added to the media at a final concentration of 8 µg/mL for one hour. Following this acute treatment, the cultures were centrifuged at 4000 × g for five minutes. Bacterial pellets were washed twice with 0.7% saline to remove MMC. Finally, bacterial pellets were suspended in fresh TSB and returned to 37°C with shaking to monitor bacterial growth, lysis, and phage induction.

#### Lysis curves

Bacterial lysis due to phage induction was evaluated by monitoring changes in culture turbidity using a FLUOstar Omega plate reader. Following MMC treatment and subsequent wash steps, bacterial cultures were dispensed at 200 μl/well into a sterile 96-well clear flat-bottom tissue culture-treated microplate (Greiner Bio-One) and incubated at 37°C with shaking. Optical density measurements at 600 nm were recorded every 30 min until stationary phase was reached. Growth measurements were repeated at least three independent times with consistent results. Data were exported and graphed using GraphPad Prism 6 software.

#### Flow cytometry

For flow cytometry, samples were collected at regular intervals spanning the initial MMC treatment to bacterial lysis. 100 μL of bacterial cultures was centrifuged at 4000 × g for 5 minutes. The resulting pellets were then suspended in 4% paraformaldehyde solution in phosphate buffer saline, pH 7.4 (PBS) (4% PFA; Biotium). Samples were fixed in this solution for 30 minutes at room temperature and then pelleted and suspended in 100 μL of sterile 0.7% saline. Samples were stored at 4°C until further analysis.

To measure “SOS” reporter activity, samples were analyzed on a Novocyte flow cytometer (Agilent). A total of 10,000 events were collected. Cells were first identified based on their forward and side scatter profile and subsequently analyzed for mNG signal. Gates to identify SOS positive events were calibrated using an untagged wild-type *E. coli* HS strain as a negative control, and an *E. coli* HS strain constitutively expressing mNG as a positive control. Data were analyzed using FlowJo software (Treestar).

Imaging flow cytometry was performed using an ImageStreamX MKII cytometer (Cytek Biosciences) using 60x magnification. Brightfield images of the bacterial cells were captured using channels 1 and 7. The 488 nm laser was used in channel 2 to visualize mNeonGreen fluorescence, and, when necessary, the 561 nm laser was used in channel 4 to capture mKate2 fluorescence. All other channels were disabled during data acquisition. A total of 10,000 events per sample were collected using the INSPIRE software. The raw image files were saved and subsequently analyzed using a custom data analysis template generated in IDEAS software version 6.2. Samples were gated based on pixel intensity variance and contrast morphology features generated with the machine learning algorithm included in IDEAS software. Gates were manually curated and adjusted by visually inspecting and excluding inappropriate events. Data from three independent experiments were exported and graphed using GraphPad Prism 6 software.

#### Live imaging

To prepare for imaging, 1 μL of either bacterial or phage samples was placed onto glass slides and covered with a glass coverslip. Routine fluorescence microscopy was performed using an Echo Revolve microscope with a 60x oil immersion objective. For super-resolution microscopy, bacterial cultures were embedded on 2.5% low melting point agarose pads and covered with #15H high-performance coverslips (Thorlabs).

For the staining of bacterial or phage DNA, 1 µL of the sample was combined with 100 µg/mL Hoechst 33342 dye (ImmunoChemistry Technologies) before mounting the samples onto a microscope slide. To selectively stain extracellular DNA, bacterial cultures were treated with the membrane impermeable DNA dye ethidium homodimer-III (EthD-III; Biotium) following manufacturer’s instructions. For membrane visualization, samples were stained with 10 µM FM4-64 (Biotium) prior to the mounting procedure.

#### Time-lapse imaging

To capture time-lapse images of Phollow phage induction, bacteria were first grown overnight in 5 mL TSB at 37°C with shaking. The next day, cultures were diluted 1:100 in fresh TSB and grown for an additional hour under identical conditions. Then, 120 μl of the bacterial culture was transferred into a well of an 8-well #1.5 high-precision slide chamber (Cellvis). To improve immobilization of bacterial cells, the chamber was previously coated with 0.1% poly-D-lysine solution (Thermo Scientific). To further create a firmly attached thin film of bacterial cells, the chamber was centrifuged at 1400 × g for 15 minutes at room temperature. Post-centrifugation, the supernatant was aspirated and non-adherent cells were gently removed by washing, three times, with 200 μL of a 0.7% saline solution. Finally, 150 μL of TSB supplemented with 0.2% low-melt agarose (Sigma) and 8 μg/mL MMC was added to the well. The chamber was left at room temperature for 5 minutes to ensure complete solidification of the agar before mounting the sample on the Zeiss Elyra 7 microscope. Imaging took place at 10-minute intervals using structured illumination.

### Super-resolution microscopy

Imaging was performed using a Zeiss Elyra 7 microscope equipped with an alpha Plan-Apochromat 63x/1.46 oil immersion lens. Unless otherwise specified, total Internal Reflection Fluorescence (TIRF) with ultra-high performance (uHP) illumination was employed. The images were captured using laser lines at 405 nm for Hoechst 33342 dye, CF405-phalloidin and mTFP, 488 nm for AausFP1, GFP and mNG, 561 nm for JF549 dye^61^ and mKate2 and 660 nm for BDP lipid stain. A Bandpass emission filter was centered at 560 nm in camera 2. The exposure time was consistently set to 25 ms, while the excitation laser power was maintained at 250 mW for all bacterial samples and 500 mW for zebrafish whole mounts. Z-stack images were acquired with the optimal slicing mode at a 0.085 µm step interval for single-point bacterial samples, leap mode at 0.125 μm intervals for bacterial time-lapses and leap mode at 0.256 μm for all zebrafish samples. For data reconstruction, Lattice 3D-SIM^2^ was utilized, scaling images to the raw data.

Super-resolution images of a whole zebrafish mount were taken with an alpha Plan-Apochromat 40x water immersion objective on a Zeiss LSM 980 Airyscan 2.0 microscope using the Airyscan SR mode. Imaging was performed using Zeiss Zen Blue v.3.2 software. To image a multi-channel z-stack image the 405 nm, 488 nm, and 561 nm lasers were used to image actin stained with CF405-phalloidin (Biotium), AausFP1virocells and DNA stained with JF549 dye^61^ (Janelia materials), respectively. The images were processed with Airyscan Deconvolution.

### Image Processing and Analysis

Maximum intensity projections, photobleaching corrections^62^ and z-depth coloring schemes were processed using FIJI (ImageJ) software^63^; three-dimensional renderings were generated with Imaris 10.1.1.

Manual creation of channel masks was necessary for images in Fig. 4a, Fig. 4c inset ‘iii’, Fig. 4d, and Fig. 4g, to account for the differences in fluorescent intensity between bacteria and phage. Additionally, masks were created in Fig. 4a to reduce spectral overlap and emphasize the unique signal of each fluorescent channel.

Analysis of intracellular viral foci was done using built-in thresholding, particle analysis, and area measurement tools in Fiji^63^. Cell lengths and number of viral foci per cell were quantified using maximum intensity projection images acquired by super-resolution microscopy. 2D Surface area measurements of viral foci were done using images of three-dimensional renderings generated in Imaris and acquired by electron microscopy.

### Transmission electron microscopy

To prepare for transmission electron microscopy (TEM), bacteria were treated with a pulse of MMC as described above. Samples were collected 1h after the MMC pulse and applied to 300- or 400-mesh carbon-coated copper grids (Gilder, Ted Pella) and negatively stained with 0.75% Uranyl Formate using established protocols^64^ and as described below. Sample concentrations were adjusted based on results from screening. Grids were negatively glow-discharged with a PELCO easiGlow (Ted Pella) to render them hydrophilic prior to staining. 3 μL of sample was applied to each grid for 2 seconds or up to 60 seconds depending on sample concentration. Then, excess liquid was blotted, the grid was washed with two drops of Milli-Q water, stained with two drops of Uranyl Formate, and allowed to air dry. For some preparations, the Milli-Q water wash step was omitted and replaced with stain. All samples were screened and data collected using a JEOL JEM 2100F or a JEOL JEM 2800 transmission electron microscope equipped with a Gatan OneView (4k x 4k) camera. Resulting micrographs were analyzed and contrast enhanced for clarity using FIJI^63^.

### Phage purification for flow virometry

Phage lytic replication was induced as described above, and 1 mL of bacterial culture was collected 1.5-2h post-wash. Chloroform (0.1x culture volume) was added and the sample was vortexed for 1 minute and incubated at room temperature for 5 minutes to lyse bacterial cells. The sample was centrifuged at 10,000xg for 5 minutes to pellet bacterial cell debris and the supernatant was moved to a new tube. 10x DNAse I buffer was added to 1x final concentration along with 1 ul of DNAse I, and the sample was incubated at 37°C for 30 minutes. EDTA (20 mM final concentration) was added to inactivate the DNAse I. NaCl (final concentration 1 M) was added and the sample was incubated on ice for 30 minutes and centrifuged at 10,000xg for 5 minutes to pellet debris. The supernatant was used for flow virometry.

### Flow virometry

Flow virometry was performed on Cytek’s Northern Lights 3-laser spectral flow cytometer or Cytek’s Aurora 5-laser spectral flow cytometer using the SpectroFlow software. Phage particle-sized events were first identified and detected on a FSC-H vs SSC-H plot using Spherotech’s 0.13 μm yellow sizing beads (NFPPS-0152-5) while adjusting voltage gains for FSC and SSC. A total of 10,000-50,000 events were captured depending on the phage availability in the samples. Raw FCS files were unmixed with reference controls of each respective Phollow phage sample containing the SpyTag and SpyCatcher, and the unstained reference control being a P2 phage deletion mutant strain.

Unmixed FCS files were analyzed using FCS Express. Phages were gated based on their size relative to the 100 μm sizing beads described above. Single particle gating on SSC-A vs SSC-H was used to discriminate against doublets. Positive MFI gates were drawn for each respective fluorescent Catcher peptide using samples negative for the SpyTag but containing the fluorescent Catcher peptide. Data were graphed and analyzed using GraphPad Prism 6 software.

### MIC determination and evaluation of phage induction with different antibiotics

To evaluate the minimum inhibitory concentration (MIC) of MMC, trimethoprim (GoldBio), and ciprofloxacin (GoldBio), Phollow virocells were first grown overnight in 5 mL TSB at 37° C with shaking. The next day, cultures were diluted 1:10^6^ in fresh TSB and grown for an additional three hours under identical conditions. Then the bacterial cultures were diluted 1:3 into fresh TSB containing a final antibiotic concentration ranging from 31.25 to 2000 ng/mL for MMC, 19.5 to 1250 ng/mL for trimethoprim, and 48 to 3125 ng/mL for ciprofloxacin. The different cultures were dispensed into a sterile 96-well clear-bottom microplate, and optical density was measured as described above. MIC was determined as the lowest concentration required to significantly impair bacterial growth without complete eradication. To evaluate the phage induction potential of the three antibiotics, the bacteria were cultured as described for the MIC and exposed to a range of antibiotic concentrations relative to the MIC, ranging from 0.05x to 2x the MIC. 130 μl of the bacteria culture was collected 24h after antibiotic exposure, and phages were purified from bacterial debris as described for virometry. To visually quantify phage induction at different antibiotic concentrations, 1 μl of purified phage samples was placed onto glass slides and covered with a glass coverslip. Fluorescence microscopy was performed using an Echo Revolve microscope with a 60x oil immersion objective. The free virions per field of view were quantified using the particle analysis function of FIJI (ImageJ)^63^. Peak virion output for each antibiotic was quantified by flow virometry as described above.

### Infectivity assay using lysogen forming units (LFU)

Phage lytic replication was induced as described above, and 500 μl of bacterial culture was collected and centrifuged at 10,000xg for 5 minutes to pellet bacterial cells. The supernatant was moved to a new tube and 0.1x volumes of chloroform was added and the sample was vortexed for 1 minute and incubated at room temperature for 5 minutes before centrifuging at 10,000xg for 5 minutes. The supernatant contained purified phage used for the infectivity assay. The recipient “target” strain (*E. coli* HS ΔP2) was diluted 1:50 in 5 mL of fresh TSB supplemented with 0.5 mM CaCl_2_ and incubated at 37°C with shaking for 30 minutes prior to infection. 100 μl of purified phage was added to the recipient strain and incubated at 37°C with shaking for 1h. Dilutions of the infection culture were plated on TSA supplemented with chloramphenicol and gentamicin to select for lysogens. Plates were incubated overnight at 37°C and colonies were counted. An *E. coli* HS *P2::clm* strain was used as a donor control to account for potential phage-independent transfer of chloramphenicol resistance during LFU assays.

### In vitro phage transmission assays

To track the horizontal transmission of Phollow phages, we used a defined two-member bacterial community consisting of a phage-donor strain and a phage-naïve target strain. The donor strain, *E. coli* HS TTW255 (HS *gpN:SpyTag_Original*; *P2 moron::clm*: *attTn7*::*sco:mNG*), is an mNG Phollow virocell that harbors a chloramphenicol resistance cassette in the second moron region of the P2 prophage. This feature facilitates the use of selective media to monitor lysogenic conversion of the target strain. The target strain, *E. coli* HS TTW269 (ΔP2; a*ttTn7*::*sco:mKate2*), has been genetically cured of the P2 prophage genome while reconstituting the *attB* attachment site to enable de novo lysogenization. Additionally, this strain constitutively expresses a SpyCatcher-mKate2 fusion protein, capable of tagging newly produced “dark” Phollow phage from lytic infection events.

Prior to the experiment, each strain was cultured independently overnight in 5 mL of TSB at 37°C with shaking. The next day, 500 μL of the saturated cultures were mixed in a microcentrifuge tube and homogenized by vortexing for 30 seconds. The resulting mixture was diluted 1:100 with fresh TSB and grown for 1h at 37°C with shaking. Subsequently, this co-culture was treated with a MMC pulse, as described above. Following the wash steps and resuspension of the pellet in TSB, the culture was further incubated at 37°C with agitation for 24h. Samples of 200 μL were taken at defined intervals: before MMC addition, immediately before the washing procedure, hourly for the subsequent 6h post-wash, and finally, 24h after the washing step. At each time point, 100 μL of the samples were treated according to the previously described imaging flow cytometry protocol. Fixed samples were stored at 4°C until further analysis.

The remaining sample volume was subjected to a ten-fold serial dilution using 0.7% saline solution. Aliquots of 4 µL from each dilution (ranging from undiluted to 10^-7^) were spot-plated onto TSA for total viable counts and onto TSA supplemented with 20 µg/mL chloramphenicol to distinguish lysogenic fractions. After an overnight incubation at 37°C, colony-forming units (CFUs) were enumerated. Differentiation between donor and target parental populations was achieved using a fluorescent stereomicroscope. For samples that did not yield countable colonies at the lowest dilution, a value of 1 was assigned to represent the limit of detection (LOD). Data from three independent experiments was graphed and analyzed using GraphPad Prism 6 software.

### In vivo characterization of phage outbreaks in the zebrafish gut

#### Induction of phage lytic replication

Germ-free zebrafish larvae were colonized with either *E. coli* HS or *Plesiomonas* AausFP1 Phollow virocells for a duration of 48h. For co-colonization experiments, a 1:1 mix of *E. coli* HS AausFP1 donor and ΔP2 mKate2 cells was used for inoculation.

To induce phage lytic replication and subsequent phage dispersal, trimethoprim was added directly into flask water containing animals at a final concentration of 125 ng/mL. Fish and flask water were sampled 4 and 24h post-treatment to evaluate bacterial abundances, virion infectivity, and to perform live imaging.

#### Harvesting bacterial and viral contents from the gut

Dissection of larval zebrafish guts was done as previously described with slight modifications^26,57^. Briefly, dissected guts of tricaine-euthanized zebrafish were harvested and placed in a 1.6 mL tube containing 500 μL sterile 0.7% saline and 100 μL 0.5 mm zirconium oxide beads (Next Advance). Guts were homogenized using a bullet blender tissue homogenizer (Next Advance) for 60 seconds on power 4.

#### Culture-based quantification of bacterial abundances

Gut and water contents were serially plated onto TSA for viable cells, or when indicated, TSA supplemented with 20 µg/mL chloramphenicol to distinguish lysogens. Plates were incubated overnight at 37°C prior to enumeration of CFUs and determination of bacterial abundances. Abundance data presented throughout the main text are pooled from a minimum of 3 independent experiments (n = 30 dissected guts and 6 waters per condition). For samples that did not yield countable colonies at the lowest dilution, a value of 1 was assigned to represent the LOD. Data were plotted and analyzed using GraphPad Prism 6 software.

#### Assessment of virion infectivity

Immediately after harvesting gut and water contents, samples were treated with 10% chloroform. These mixtures were vigorously homogenized for 30 seconds using a vortex. Subsequently, the solutions were incubated at room temperature for 10 minutes before centrifugation at 10000 × g for 3 minutes. The aqueous solutions containing purified virions were carefully extracted and further used in infectivity assays.

To infect an *E. coli* ΔP2 mKate2 target strain, an overnight culture of the strain was diluted 1:100 in fresh TSB and incubated at 37°C with shaking for 30 minutes. The culture was then dispensed in 100 µL aliquots into the wells of a deep-well 96-well plate (Fisher Scientific). Centrifugation of the plate was performed at 4000 × g for 5 minutes to pellet the cells. The pellets were washed twice with one volume of 0.7% saline solution.

Finally, the cell pellets were suspended in 100 µL of the chloroform-treated solutions. The suspensions were incubated at 37°C with agitation for one hour. Following the incubation, the cultures were serially diluted and plated onto TSA supplemented with chloramphenicol to select for de novo lysogens. The plates were incubated overnight at 37°C, after which the LFUs were enumerated. For samples that did not yield countable colonies at the lowest dilution, a value of 1 was assigned to represent the LOD. Data from 3 independent experiments were graphed and analyzed using GraphPad Prism 6 software. Statistical analysis was performed using one-way ANOVA with a Kruskal-Wallis and Dunn’s multiple comparisons test.

#### Microscopy

Zebrafish larvae were fixed in 4% PFA for two hours, after which they were washed twice with PBS. Subsequently, up to five larvae were placed into a microcentrifuge tube and resuspended in 200 µL of PBS containing 1 µM JF549 dye^61^ for nuclear DNA staining and 1 unit of CF405-phalloidin for actin filament staining. For experiments involving TgBAC(*nkx2.2a*:*megfp)* reporter fish or those displaying two-member bacterial communities, Hoechst 33342 DNA dye was utilized instead at a concentration of 2 µg/mL. The larvae remained in the staining solution overnight or until ready for imaging. When indicated, stained animals were incubated with 0.5 µg/mL 650/665 BDP (Luminiprobe) overnight to stain for lipid droplets. To prepare for imaging, each specimen was positioned onto a microscope slide, ensuring the removal of all excess liquid, followed by the application of 5 µL of EverBrite mounting media (Biotium). For dissected organs, 3% methylcellulose was used as mounting media instead. Finally, a coverslip was placed over the larvae. Any residual mounting media was carefully dried out, and the coverslip was securely sealed with nail polish.

For live imaging experiments, larvae were anesthetized with 0.4% Tricaine to minimize movement and were then embedded in 1% low-melting-point agarose placed in 35 mm glass-bottom Petri dishes. The plate was finally filled with sterile EM.

For the super-resolution imaging of a whole animal, fixed and stained larvae were embedded in 1% low-melting-point agarose placed in 35 mm glass-bottom Petri. After agarose solidification, 3 mL of deionized water was added to the plate to reduce evaporation.

Images were acquired as described in the “Super-resolution microscopy” section.

---

**Extended Data Table 1.**
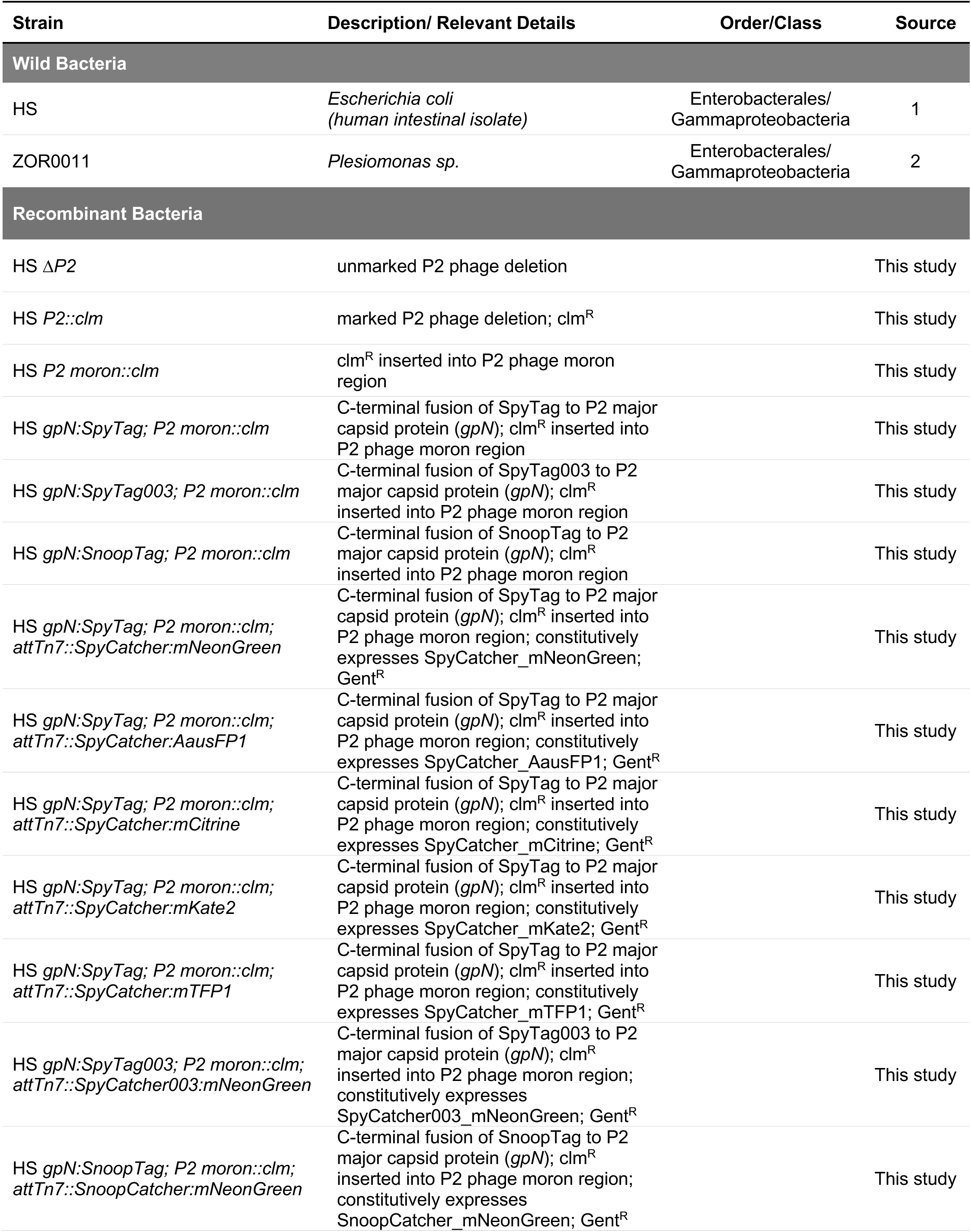
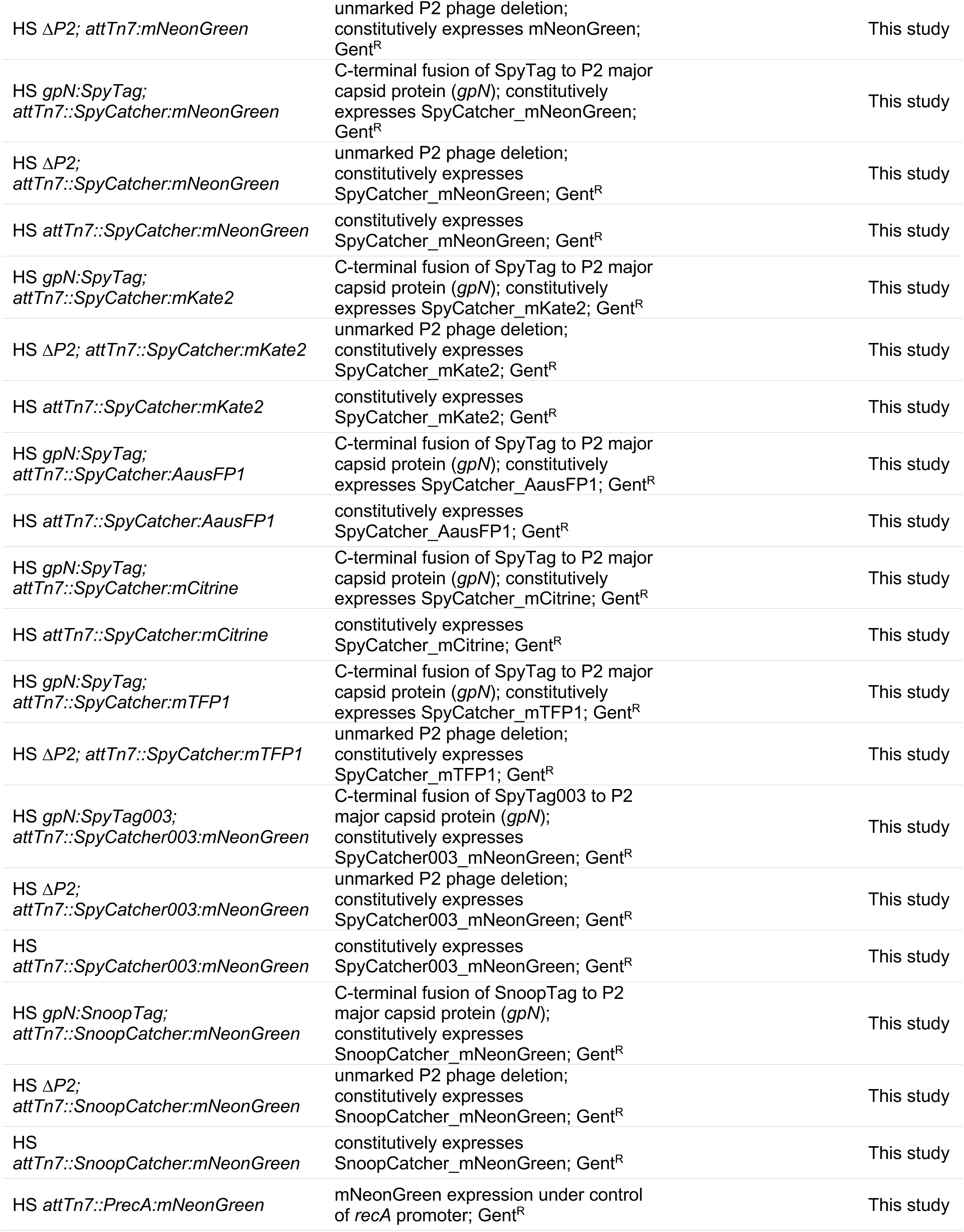
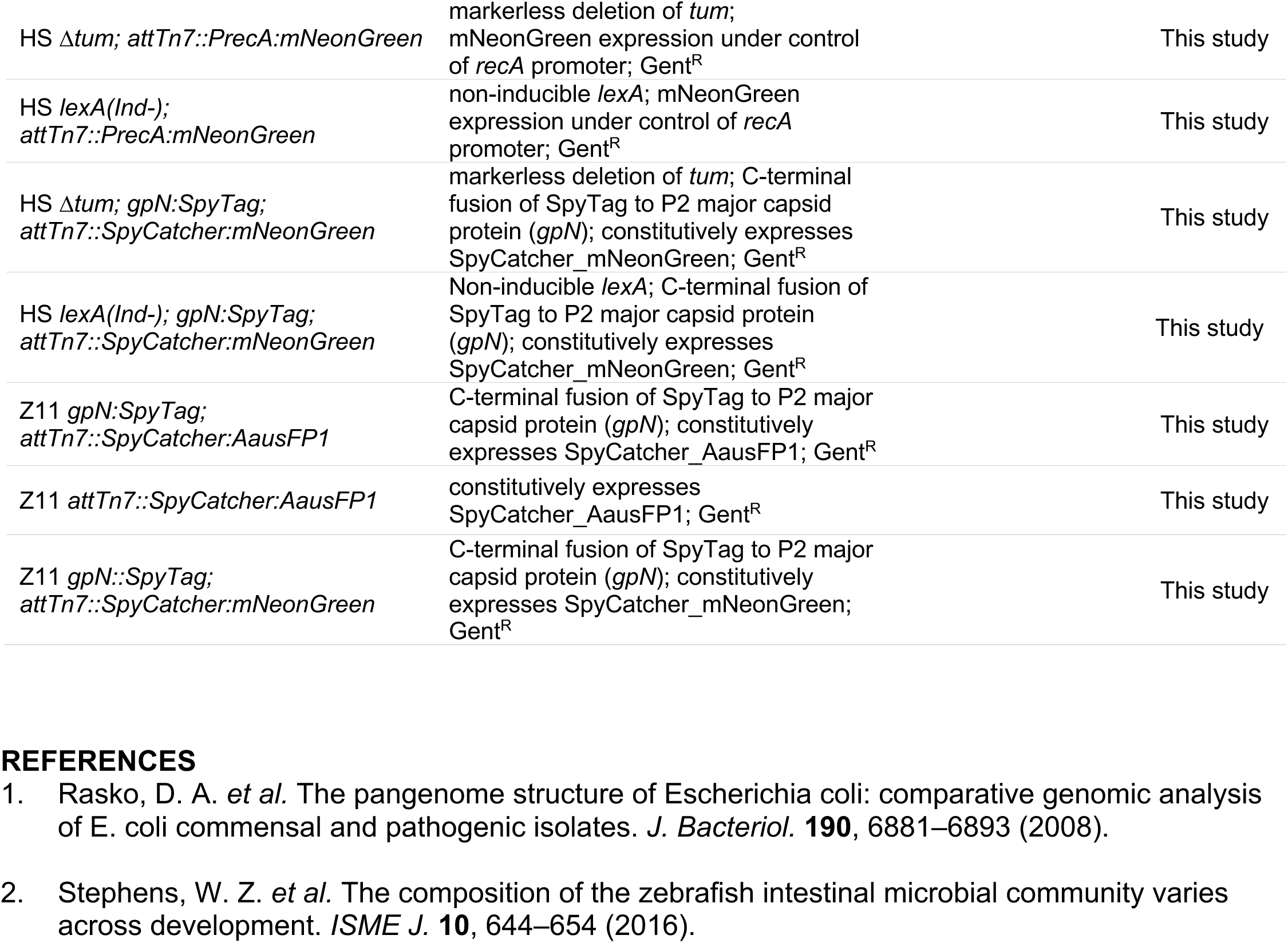
Wild and Recombinant Bacteria.

| Strain | Description/ Relevant Details | Order/Class | Source |
| --- | --- | --- | --- |
| **Wild Bacteria** |  |  |  |
| HS | *Escherichia coli*<br>(human intestinal isolate) | Enterobacterales/<br>Gammaproteobacteria | 1 |
| ZOR0011 | *Plesiomonas* sp. | Enterobacterales/<br>Gammaproteobacteria | 2 |
| **Recombinant Bacteria** |  |  |  |
| HS ΔP2 | unmarked P2 phage deletion | | This study |
| HS P2::clm | marked P2 phage deletion; clm<sup>R</sup> |  | This study |
| HS P2 moron::clm | clm<sup>R</sup> inserted into P2 phage moron region |  | This study |
| HS gpN:SpyTag; P2 moron::clm | C-terminal fusion of SpyTag to P2 major capsid protein (gpN); clm<sup>R</sup> inserted into P2 phage moron region |  | This study |
| HS gpN:SpyTag003; P2 moron::clm | C-terminal fusion of SpyTag003 to P2 major capsid protein (gpN); clm<sup>R</sup> inserted into P2 phage moron region |  | This study |
| HS gpN:SnoopTag; P2 moron::clm | C-terminal fusion of SnoopTag to P2 major capsid protein (gpN); clm<sup>R</sup> inserted into P2 phage moron region |  | This study |
| HS gpN:SpyTag; P2 moron::clm;<br>attTn7::SpyCatcher:mNeonGreen | C-terminal fusion of SpyTag to P2 major capsid protein (gpN); clm<sup>R</sup> inserted into P2 phage moron region; constitutively expresses SpyCatcher_mNeonGreen; Gent<sup>R</sup> |  | This study |
| HS gpN:SpyTag; P2 moron::clm;<br>attTn7::SpyCatcher:AausFP1 | C-terminal fusion of SpyTag to P2 major capsid protein (gpN); clm<sup>R</sup> inserted into P2 phage moron region; constitutively expresses SpyCatcher_AausFP1; Gent<sup>R</sup> |  | This study |
| HS gpN:SpyTag; P2 moron::clm;<br>attTn7::SpyCatcher:mCitrine | C-terminal fusion of SpyTag to P2 major capsid protein (gpN); clm<sup>R</sup> inserted into P2 phage moron region; constitutively expresses SpyCatcher_mCitrine; Gent<sup>R</sup> |  | This study |
| HS gpN:SpyTag; P2 moron::clm;<br>attTn7::SpyCatcher:mKate2 | C-terminal fusion of SpyTag to P2 major capsid protein (gpN); clm<sup>R</sup> inserted into P2 phage moron region; constitutively expresses SpyCatcher_mKate2; Gent<sup>R</sup> |  | This study |
| HS gpN:SpyTag; P2 moron::clm;<br>attTn7::SpyCatcher:mTFP1 | C-terminal fusion of SpyTag to P2 major capsid protein (gpN); clm<sup>R</sup> inserted into P2 phage moron region; constitutively expresses SpyCatcher_mTFP1; Gent<sup>R</sup> |  | This study |
| HS gpN:SpyTag003; P2 moron::clm;<br>attTn7::SpyCatcher003:mNeonGreen | C-terminal fusion of SpyTag003 to P2 major capsid protein (gpN); clm<sup>R</sup> inserted into P2 phage moron region; constitutively expresses SpyCatcher003_mNeonGreen; Gent<sup>R</sup> |  | This study |
| HS gpN:SnoopTag; P2 moron::clm;<br>attTn7::SnoopCatcher:mNeonGreen | C-terminal fusion of SnoopTag to P2 major capsid protein (gpN); clm<sup>R</sup> inserted into P2 phage moron region; constitutively expresses SnoopCatcher_mNeonGreen; Gent<sup>R</sup> |  | This study |

|  |  |  |
| --- | --- | --- |
| HS $\Delta P2$; *attTn7::mNeonGreen* | unmarked P2 phage deletion;<br>constitutively expresses mNeonGreen;<br>Gent<sup>R</sup> | This study |
| HS *gpN::SpyTag*;<br>*attTn7::SpyCatcher::mNeonGreen* | C-terminal fusion of SpyTag to P2 major<br>capsid protein (*gpN*); constitutively<br>expresses SpyCatcher_mNeonGreen;<br>Gent<sup>R</sup> | This study |
| HS $\Delta P2$;<br>*attTn7::SpyCatcher::mNeonGreen* | unmarked P2 phage deletion;<br>constitutively expresses<br>SpyCatcher_mNeonGreen; Gent<sup>R</sup> | This study |
| HS *attTn7::SpyCatcher::mNeonGreen* | constitutively expresses<br>SpyCatcher_mNeonGreen; Gent<sup>R</sup> | This study |
| HS *gpN::SpyTag*;<br>*attTn7::SpyCatcher::mKate2* | C-terminal fusion of SpyTag to P2 major<br>capsid protein (*gpN*); constitutively<br>expresses SpyCatcher_mKate2; Gent<sup>R</sup> | This study |
| HS $\Delta P2$; *attTn7::SpyCatcher::mKate2* | unmarked P2 phage deletion;<br>constitutively expresses<br>SpyCatcher_mKate2; Gent<sup>R</sup> | This study |
| HS *attTn7::SpyCatcher::mKate2* | constitutively expresses<br>SpyCatcher_mKate2; Gent<sup>R</sup> | This study |
| HS *gpN::SpyTag*;<br>*attTn7::SpyCatcher::AausFP1* | C-terminal fusion of SpyTag to P2 major<br>capsid protein (*gpN*); constitutively<br>expresses SpyCatcher_AausFP1; Gent<sup>R</sup> | This study |
| HS *attTn7::SpyCatcher::AausFP1* | constitutively expresses<br>SpyCatcher_AausFP1; Gent<sup>R</sup> | This study |
| HS *gpN::SpyTag*;<br>*attTn7::SpyCatcher::mCitrine* | C-terminal fusion of SpyTag to P2 major<br>capsid protein (*gpN*); constitutively<br>expresses SpyCatcher_mCitrine; Gent<sup>R</sup> | This study |
| HS *attTn7::SpyCatcher::mCitrine* | constitutively expresses<br>SpyCatcher_mCitrine; Gent<sup>R</sup> | This study |
| HS *gpN::SpyTag*;<br>*attTn7::SpyCatcher::mTFP1* | C-terminal fusion of SpyTag to P2 major<br>capsid protein (*gpN*); constitutively<br>expresses SpyCatcher_mTFP1; Gent<sup>R</sup> | This study |
| HS $\Delta P2$; *attTn7::SpyCatcher::mTFP1* | unmarked P2 phage deletion;<br>constitutively expresses<br>SpyCatcher_mTFP1; Gent<sup>R</sup> | This study |
| HS *gpN::SpyTag003*;<br>*attTn7::SpyCatcher003::mNeonGreen* | C-terminal fusion of SpyTag003 to P2<br>major capsid protein (*gpN*);<br>constitutively expresses<br>SpyCatcher003_mNeonGreen; Gent<sup>R</sup> | This study |
| HS $\Delta P2$;<br>*attTn7::SpyCatcher003::mNeonGreen* | unmarked P2 phage deletion;<br>constitutively expresses<br>SpyCatcher003_mNeonGreen; Gent<sup>R</sup> | This study |
| HS<br>*attTn7::SpyCatcher003::mNeonGreen* | constitutively expresses<br>SpyCatcher003_mNeonGreen; Gent<sup>R</sup> | This study |
| HS *gpN::SnoopTag*;<br>*attTn7::SnoopCatcher::mNeonGreen* | C-terminal fusion of SnoopTag to P2<br>major capsid protein (*gpN*);<br>constitutively expresses<br>SnoopCatcher_mNeonGreen; Gent<sup>R</sup> | This study |
| HS $\Delta P2$;<br>*attTn7::SnoopCatcher::mNeonGreen* | unmarked P2 phage deletion;<br>constitutively expresses<br>SnoopCatcher_mNeonGreen; Gent<sup>R</sup> | This study |
| HS<br>*attTn7::SnoopCatcher::mNeonGreen* | constitutively expresses<br>SnoopCatcher_mNeonGreen; Gent<sup>R</sup> | This study |
| HS *attTn7::PrecA::mNeonGreen* | mNeonGreen expression under control<br>of *recA* promoter; Gent<sup>R</sup> | This study |
| HS $\Delta tum$; *attTn7::PrecA:mNeonGreen* | markerless deletion of *tum*; mNeonGreen expression under control of *recA* promoter; Gent<sup>R</sup> | This study |
| HS *lexA(Ind-)*; *attTn7::PrecA:mNeonGreen* | non-inducible *lexA*; mNeonGreen expression under control of *recA* promoter; Gent<sup>R</sup> | This study |
| HS $\Delta tum$; *gpN::SpyTag*; *attTn7::SpyCatcher:mNeonGreen* | markerless deletion of *tum*; C-terminal fusion of SpyTag to P2 major capsid protein (*gpN*); constitutively expresses SpyCatcher_mNeonGreen; Gent<sup>R</sup> | This study |
| HS *lexA(Ind-)*; *gpN::SpyTag*; *attTn7::SpyCatcher:mNeonGreen* | Non-inducible *lexA*; C-terminal fusion of SpyTag to P2 major capsid protein (*gpN*); constitutively expresses SpyCatcher_mNeonGreen; Gent<sup>R</sup> | This study |
| Z11 *gpN::SpyTag*; *attTn7::SpyCatcher:AausFP1* | C-terminal fusion of SpyTag to P2 major capsid protein (*gpN*); constitutively expresses SpyCatcher_AausFP1; Gent<sup>R</sup> | This study |
| Z11 *attTn7::SpyCatcher:AausFP1* | constitutively expresses SpyCatcher_AausFP1; Gent<sup>R</sup> | This study |
| Z11 *gpN::SpyTag*; *attTn7::SpyCatcher:mNeonGreen* | C-terminal fusion of SpyTag to P2 major capsid protein (*gpN*); constitutively expresses SpyCatcher_mNeonGreen; Gent<sup>R</sup> | This study |

**Extended Data Table 2.** Summary of observations for Phollow phage tagging systems.

| **Description** | **Snoop** | **Spy003** | **SpyO** |
| --- | --- | --- | --- |
| Effect in bacterial growth<br>And morphology | Pronounced lysis. Altered bacterial morphology. | Growth and lysis are similar to wild-type strain. | Growth and lysis are similar to wild-type strain. |
| Phage morphology and dynamics | Small foci, thin ribbon structures. Delayed virion detection compared to SpyTags | Large foci, long ribbon and serpentine structures. | Large foci, long ribbon and serpentine structures. |
| Microscopy of virions | Small, dim and sparse particles | Large, bright and abundant particles. | Large and bright particles. |
| Flow virometry | Strong detection with an average of 58.51% positive events. | Moderate detection with an average of 33.95% positive events. | Strong detection with an average 49.12% positive events. |
| Infectivity | Similar to wild-type strain. | Similar to wild-type strain. | Similar to wild-type strain. |

**Extended Data Table 3.**
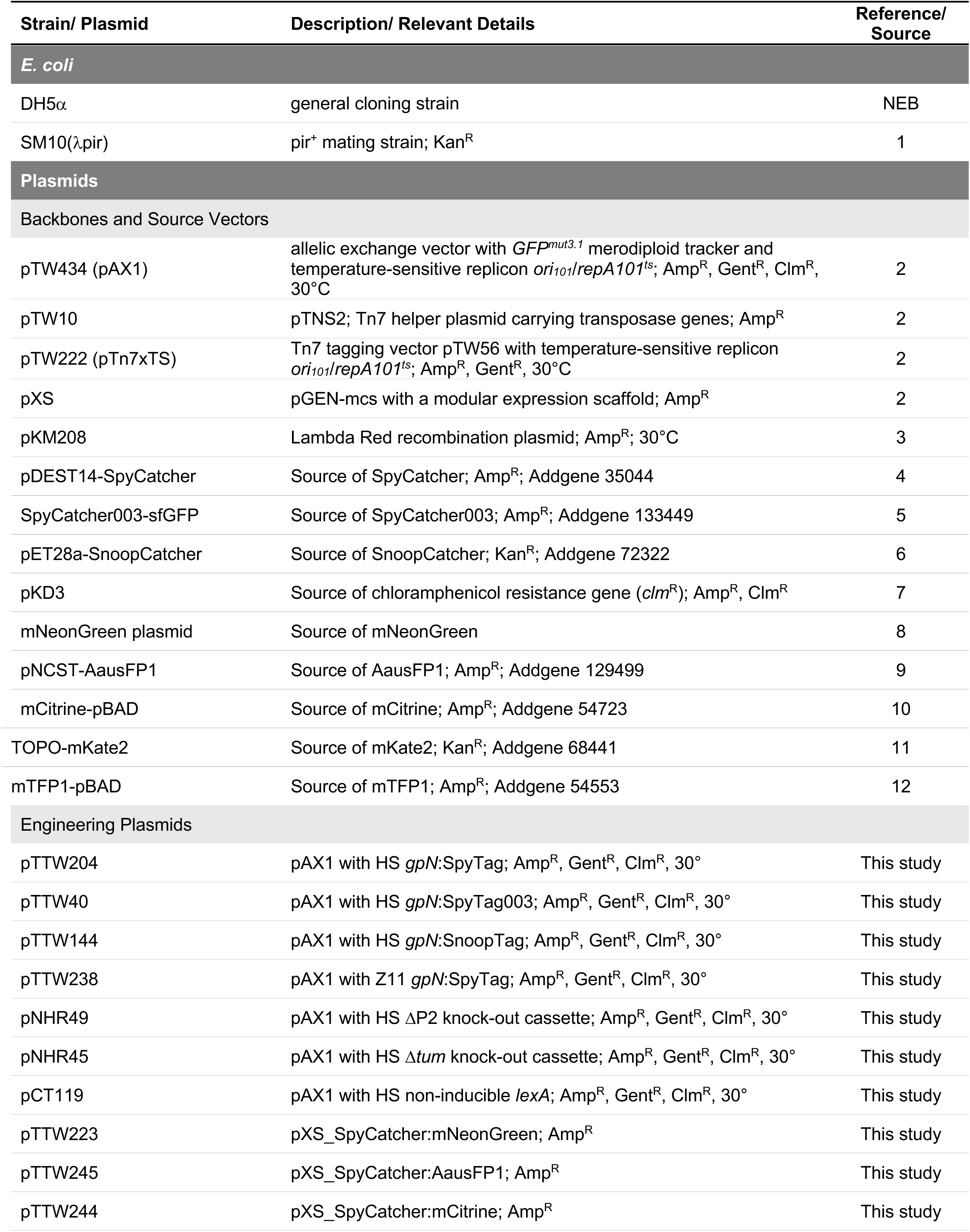
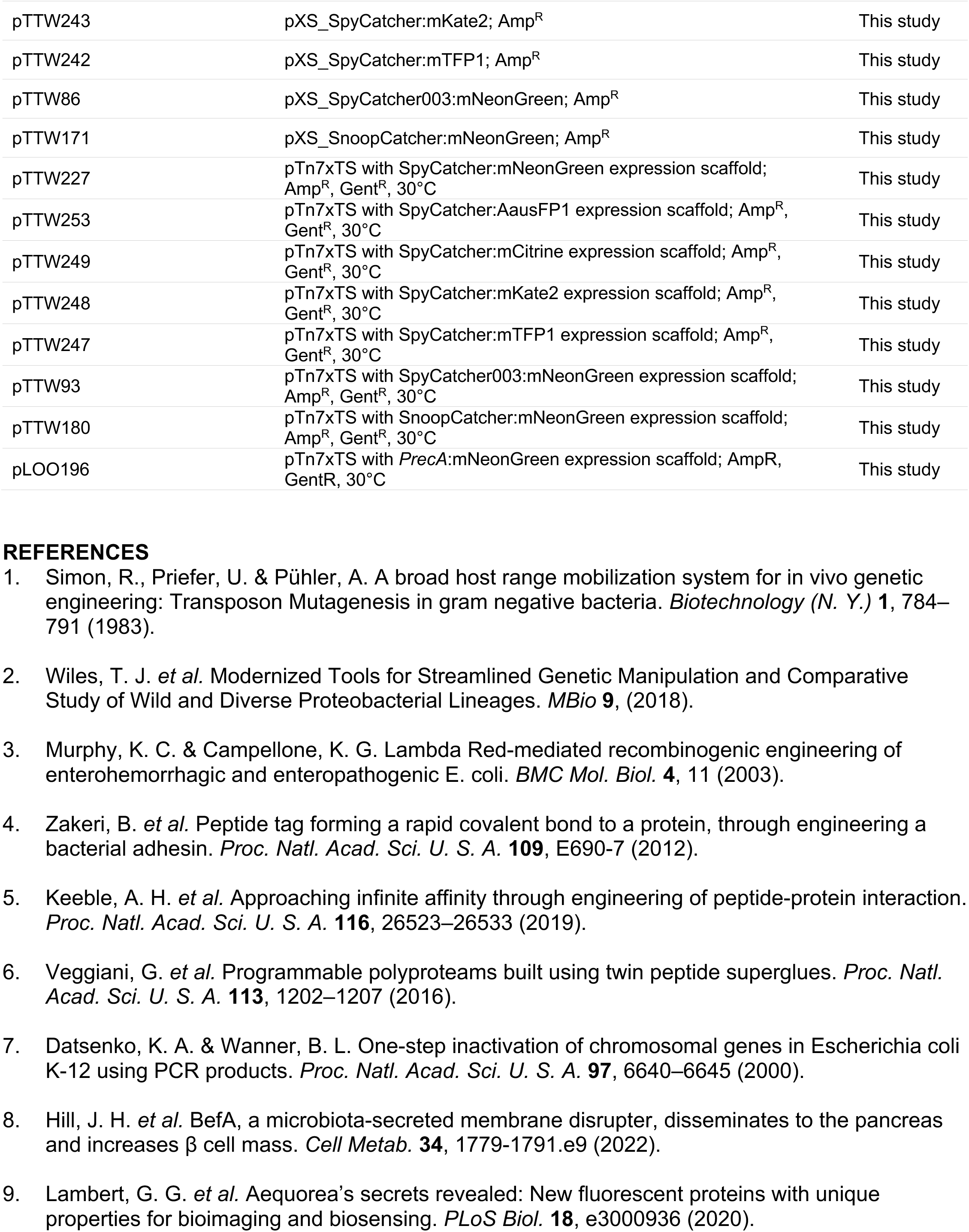
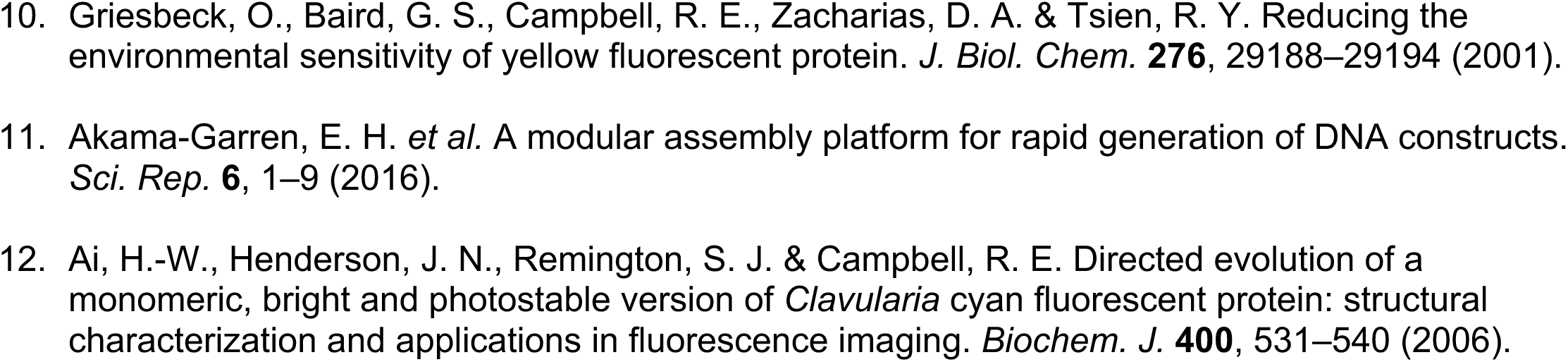
*E. coli* Strains and Plasmids.

**Extended Data Table 4.**
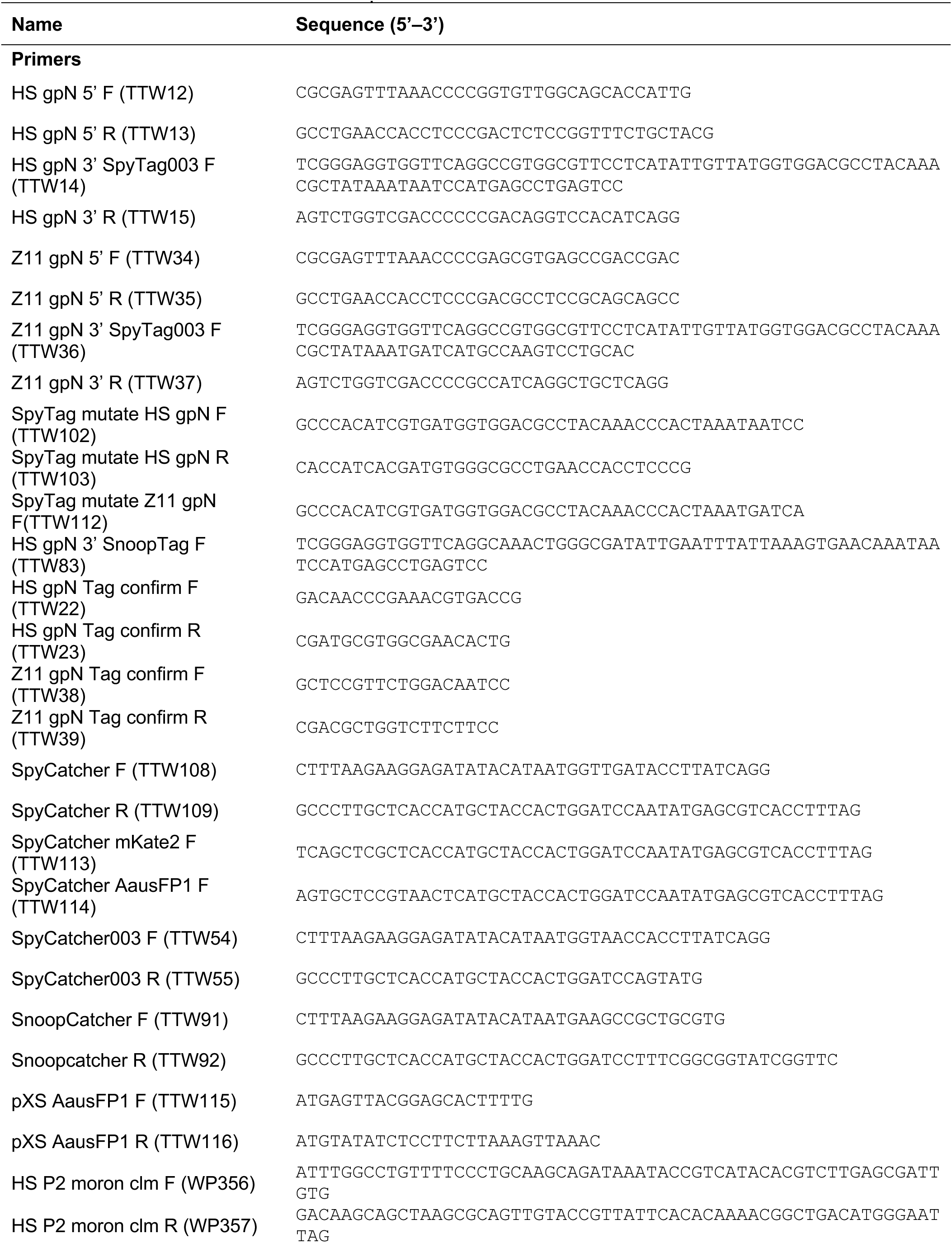
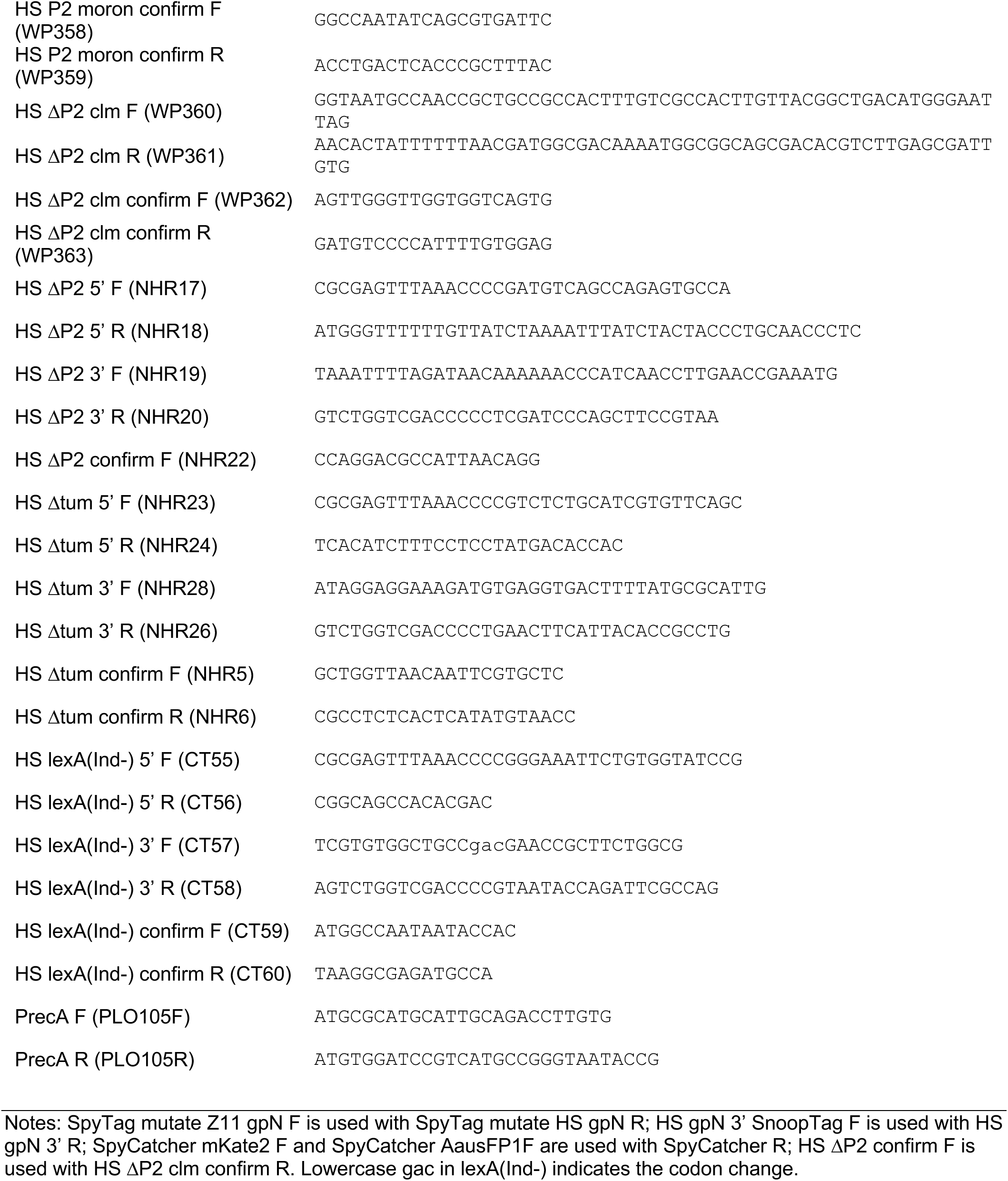
Primer sequences.

## Supplementary File 1. Plasmid Construction.

*Note: When referring to specific genetic loci within bacterial chromosomes, locus tags are provided that can be used to retrieve sequence information from the National Center for Biotechnology Information (NCBI) website (*https://www.ncbi.nlm.nih.gov/).

### C-terminal fusion of tags to the major capsid protein

An allelic exchange cassette was constructed by amplifying the 5’ homology region (846 bp; primers TTW12&13) just upstream of the HS *gpN* stop codon (locus tag ECHS_RS04530) and the 3’ homology region (837 bp; primers TTW14&15) which included the *gpN* stop codon and downstream region. The DNA sequence for a six amino acid linker (SGGGSG) and the appropriate tag (SpyTag003 or SnoopTag) were incorporated into the forward primer for the 3’ homology region just upstream of the stop codon. The two homology arms were inserted into the SmaI linearized pAX1 vector using in vivo assembly (IVA). The same strategy, amplifying a 5’ homology region (810 bp; primers TTW34&35) and 3’ homology region (793 bp; primers TTW36&37), was used to tag the ZOR0011 major capsid protein *gpN* (locus tag L975_RS01430). Site directed mutagenesis via IVA was used to convert pAX1 *gpN*:SpyTag003 to pAX1 *gpN*:SpyTag for both HS (primers TTW102&103) and ZOR0011 (primers TTW103&112).

### Deletion of P2 prophage

An allelic exchange cassette was constructed to create a markerless deletion of the P2 phage leaving the *attB* site intact on the bacterial chromosome. The genomic regions upstream (956 bp; primers NHR17&18; starting with *rcdA*; locus tag ECHS_RS04440) and downstream (1001 bp; primers NHR19&20; starting 77bp downstream of *ogr*; locus tag ECHS_RS04645) of the P2 phage were amplified such that the resulting plasmid would delete the P2 phage and recreate the *attB* site. The two homology arms were inserted into the SmaI linearized pAX1 vector using IVA.

### Deletion of *tum*

An allelic exchange cassette was constructed to create a markerless deletion of *tum* (locus tag ECHS_RS04495). The genomic regions upstream (969 bp; primers NHR23&24) and downstream (859 bp; primers NHR26&28) of the *tum* gene were amplified such that the resulting plasmid would delete the gene and fuse the start and stop codons. The two homology arms were inserted into the SmaI linearized pAX1 vector using IVA.

### Non-inducible *lexA* (*lexA*(Ind-))

An allelic exchange cassette was constructed to create a markerless G85D mutation of *lexA* (locus tag ECHS_RS21095). The genomic regions upstream (841bp; primers CT55&56) and downstream (888 bp; CT57&58) of the modified codon were amplified such that the resulting plasmid would replace the native GGT nucleotide sequence with GAC. The two homology arms were inserted into the SmaI linearized pAX1 vector using IVA.

### N-terminal fusion of Catcher proteins to fluorescent proteins

The catcher sequence was amplified from the appropriate plasmid: SpyCatcher (pDEST14-SpyCatcher was a gift from Mark Howarth; Addgene plasmid # 35044) Zakeri et al., SpyCatcher003 (SpyCatcher003-sfGFP was a gift from Mark Howarth; Addgene plasmid # 133449) Keeble et al., SnoopCatcher (pET28a-SnoopCatcher was a gift from Mark Howarth; Addgene plasmid # 72322) Veggiani et al., and inserted into pXS linearized with NdeI and containing a gene for a fluorescent protein (mNeonGreen, mCitrine, mKate2, mTFP1) using IVA. Because AausFP1 contains two NdeI sites, pXS_AausFP1 was amplified by PCR to generate a linear plasmid for IVA. The complete cassette containing a *Ptac* promoter, Catcher sequence, fluorescent protein sequence and transcriptional terminator was subcloned from pXS with EcoRI and AvrII and ligated into SmaI linearized pTn7xTS.

### Fluorescent SOS reporter

A fluorescent SOS reporter was engineered by amplifying the 120 bp promoter region upstream of the *E. coli* HS *recA* gene (locus tag ECHS_RS13985, P*recA*) using primers PrecA_F and PrecA_R. Engineered restriction sites flanking the amplicon (SphI and BamHI) were then used to insert this fragment into a variant of the pTn7xTS delivery vector harboring an open reading frame encoding mNeonGreen (mNG) with an epsilon enhancer and a consensus Shine-Dalgarno sequence within the 5’ untranslated region (UTR), as well as the synthetic transcriptional terminator L3S2P21 in the 3’ UTR. The reporter was inserted into the *attTn7* site of the bacterial chromosome via Tn7 tagging.

